# Speed and correctness guarantees for programmable enthalpy-neutral DNA reactions

**DOI:** 10.1101/2022.04.13.488226

**Authors:** Boya Wang, Chris Thachuk, David Soloveichik

**Author notes:** Preliminary versions of some of the results appeared in conference proceedings [1].

## Abstract

Molecular control circuits embedded within chemical systems to direct molecular events have transformative applications in synthetic biology, medicine, and other fields. However, it is challenging to understand the collective behavior of components due to the combinatorial complexity of possible interactions. Some of the largest engineered molecular systems to date have been constructed from DNA strand displacement reactions, in which signals can be propagated without a net change in base pairs. For linear chains of such enthalpy-neutral displacement reactions, we develop a rigorous framework to reason about interactions between regions that must be complementary. We then analyze desired and undesired properties affecting speed and correctness of such systems, including the spurious release of output (leak) and reversible unproductive binding (toehold occlusion), and experimentally confirm the predictions. Our approach, analogous to the rigorous proofs of algorithm correctness in computer science, can guide engineering of robust and efficient molecular algorithms.

---

Nature evolved sophisticated molecular systems with diverse functional behaviors, ranging from simple chemical switches to vast gene regulatory networks. Inspired by nature, artificial molecular systems have shown significant promise in approaching this level of complexity. Guided by ideas from computer science and electrical engineering, molecular systems constructed from first principles are finding applications in chemistry, material sciences and medicine. Cascades of DNA strand displacement reactions have been shown to be a powerful mechanism for engineering molecular information processing and dynamics [2]. In strand displacement cascades, a DNA strand displaces another strand from multi-stranded pre-hybridized complex, and the displaced strand can in turn act as a displacing strand for downstream reactions [3]. Strand displacement cascades have yielded some of the largest biochemical systems designed from scratch [4].

Many desired and undesired properties of strand displacement systems are expressed and understood in terms of arguments about enthalpy and entropy, where the first is shorthand for the number of bonds and the second for the number of separate complexes. For example, an important obstacle to scaling up strand displacement systems is *leak*, which occurs when undesired reactions get spontaneously triggered in the absence of the correct input. A common approach to lowering leak involves adding one to three nucleotides (a *clamp*) at the end of helices which must break in the process of leak, leading to an enthalpic barrier [5, 6, 4, 7, 8]. Since leak results from a spurious interaction between different complexes, lowering their concentration increases the entropic barrier to leak. In typical strand displacement systems, leak can result from a spurious interaction between just two complexes. In a recent leak reduction method (“*NLD* design”) [9, 10], the entropy penalty to leak can be programmed to be higher. Physically separating reaction species that may react spuriously also increases the entropic barrier to leak [11]. Other undesired properties include slowing of the desired reaction pathways due to the temporary sequestration of complexes in unreactive states (“toehold occlusion” and “spurious displacement”, see below). Almost universally, complexes have parts that are complementary by design, and thus they are driven to bind together by enthalpy, making some amount of sequestration unavoidable.

The driving force for the desired strand displacement reactions is often enthalpic—the formation of 5–7 new base pairs called a toehold. However, a type of strand displacement called *toehold-exchange* can preserve the overall number of base pairs and is driven by entropy alone [12]. Such enthalpy-neutral strand displacement cascades have played an important role in *de novo* engineering molecular systems. Driven forward solely by the formation of additional separate complexes, molecular signals can get exponentially amplified [13]. In the form of “see-saw gates”, enthalpy-neutral strand displacement was a key module in chemical circuits for performing square-root computation [4], and neural network computation [14, 15]. Molecular robots that can move along a track and sort cargo [16], synthetic distributed algorithms [17], and dynamical systems including oscillators [8], have relied in part or in whole on enthalpy-neutral strand displacement modules. Since mismatches between the invading strand and its target can significantly affect the equilibrium of enthalpy-neutral displacement, single-nucleotide mismatches can be detected with high specificity [18], giving rise to potential medical applications [19, 20].

Even the simplest kind of enthalpy-neutral strand displacement cascades—a cascade of translators, which are logically equivalent to repeater gates—can already exhibit complex and useful behavior. Translators can also be composed to compute the logic *OR* of many distinct inputs translated to a common output [10]. A catalytic system can be constructed if a translator chain’ s output is the same as its input. Indeed, the powerful “see-saw” motif discussed above is a version of the enthalpy neutral translator.

Here we demonstrate an approach to systematically understand desired and undesired properties of strand displacement cascades based on combinatorial arguments about domain-level structure, enthalpy and entropy. We focus on cascades of enthalpy-neural (*EN*) translators, and show that the entire design space can be captured by adjusting two parameters describing the length of the double-stranded region of the complexes (defined by *N*) and the length of the sequence that overlaps between two logically consecutive complexes (defined by *shift*). Each of the variety of designs achieved has unique properties. Some parameter choices lead to toehold occlusion [4, 8] (blocking the toehold for a desired reaction), and spurious strand displacement (partial displacement of a strand on a complex by a spurious invader), both of which are likely to hinder the kinetics of the intended reaction pathway. We further develop combinatorial arguments about entropy and enthalpy that lead to a general analysis of leak in *EN* cascades. When comparing two configurations, if the number of bonds is the same while one configuration has *n* fewer separate complexes than the other, then the first configuration is considered unfavorable as it incurs *n* units of entropic penalty relative to the second (“One unit of entropic penalty” is approximately the energy of binding 7 base pairs; Section S1). Similarly, all else being equal, a configuration with *n* fewer bonds than another configuration is unfavorable and has a relative enthalpic penalty of *n* units. In these terms, we prove that in *EN* cascades, there is a thermodynamic penalty to leak consisting of an enthalpy penalty of 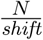, combined with an entropy penalty of 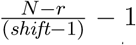even if *r* bonds break. Compared to the *NLD* design, enthalpy-neutral cascades are expected to have significantly less leak under the same conditions (see Section S8).

In an enthalpy-neutral cascade, every reaction is reversible, and thus the signal tends to spread out across the cascade more than in designs driven enthalpically by the bonding of toeholds. However, we show that the spreading does not limit the capability of enthalpy-neutral cascades to effectively propagate signal. We prove that the system completion level does not decrease arbitrarily with the depth of the cascade, allowing long cascades to be constructed. Since the leak penalty increases with larger *N* (keeping *shift* fixed), while the level of desired output is bounded from below, the “signal-to-noise” ratio can be made arbitrarily high. Because leak is reduced even at high concentrations, complex *EN* cascades may take minutes to complete instead of hours as is current state of the art [4].

We conclude with experimental demonstrations of *EN* designs with different sets of parameters. The laboratory results confirm our theoretical categorization of designs with regard to leak and undesirable kinetic properties of toehold occlusion and spurious displacement, as well as the desired signal propagation. The leak reduction we observe compares favorably with the previously proposed *NLD* designs, in terms of the kinetic leak rate, the relative triggered signal to leak ratio and the maximum amount of leak generated. Despite the inevitable complexity of interactions between complementary regions in strand displacement cascades, our work suggests that this complexity can be effectively understood and programmed.

## Design space of enthalpy-neutral strand displacement cascades

### System description

In toehold exchange (Figure 1a), a DNA strand first binds the *toehold*—an unbound single-stranded region (usually 5 to 8 nt) of the pre-hybridized complex. Subsequently, via a random walk process (3-way branch migration), this strand competes with the originally bound one, with complete displacement occurring after the dissociation of a symmetric toehold-size region. Single-stranded DNA fulfills the role of *signals* that carry information, while pre-hybridized DNA complexes (*fuels*) provide the material for the output signal.

**Figure 1:**
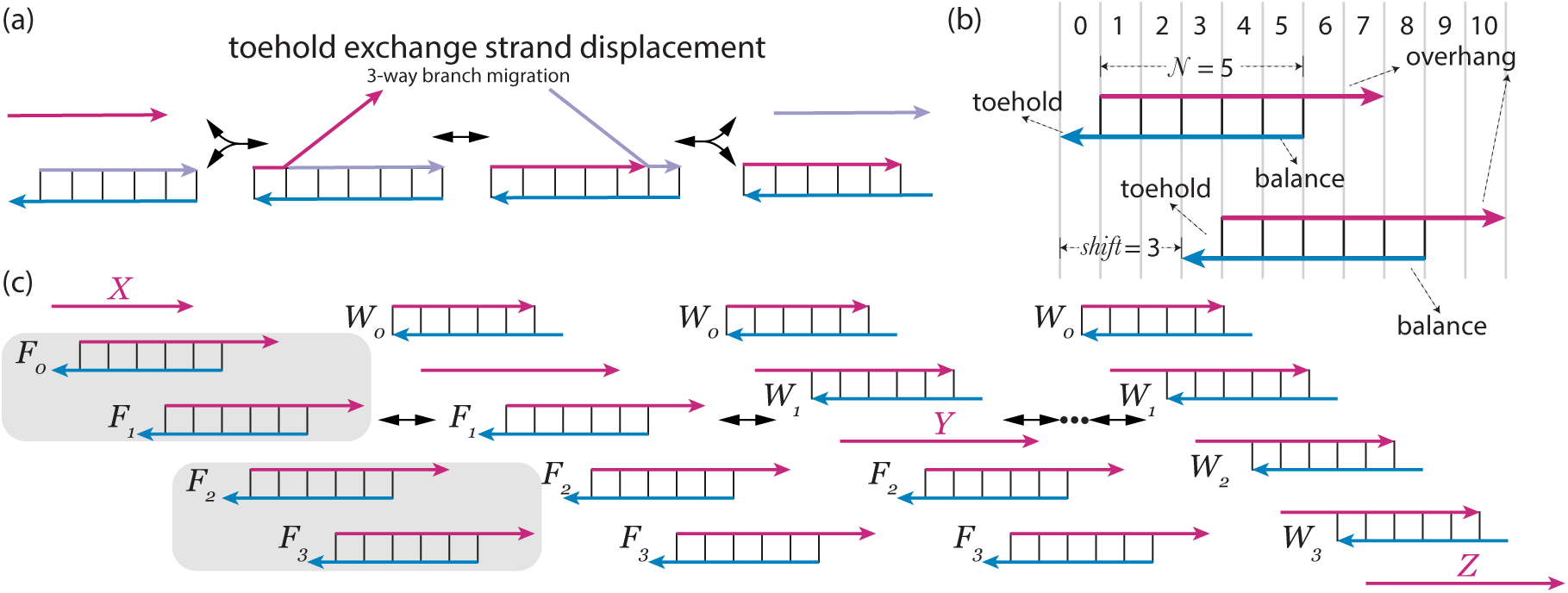
(a) The fundamental reaction step we consider. (b) The conventions of the *EN* design. (c) Desired reaction pathway of a 2-translator cascade (*X* → *Y* → *Z*). After 4 strand displacement steps, the signal strand *X* is translated to signal strand *Z*.

We use *top strand* /*bottom strand* to indicate the strand at the top/bottom of a fuel complex respectively in our illustrations. Strands are divided into domains (a unit of concatenated DNA bases). Here all domains are assumed to be short enough that strands bound by a single domain can dissociate (see Figure 1a); such domains are typically said to be of toehold-length). Strands bound by two or more domains cannot dissociate. We assume that all domains are orthogonal (no cross-talk, see discussion in Section S9).

Multiple single-stranded top domains on a fuel together are called an *overhang*. Single-stranded bottom domains are called *toeholds*. The domains at the right end of a double-stranded helix are called *balance domains*, since they break after strand displacement ensuring equal no net enthalpy change. (The *balance domain* can also be thought of as a long “clamp” in previous strand displacement designs used to “clamp-down” the ends of helices to reduce leak.) In our illustrations, domains aligned vertically have the same or complementary sequence, which is shown using *position* numbering as on top of Figure 1b. Note that as is typical of at-scale strand displacement experiments, there are many copies of each fuel although only one copy is drawn.

### Linear strand displacement cascades

When the input signal strand is present, it reacts with the first fuel displacing the top strand, which then triggers the downstream fuel (Figure 1c). A linear strand displacement cascade can be used to build *translators*, converting an input signal strand to a sequence-independent output signal strand. Figure 1c shows two translators of two fuels each (*X, Y* and *Z* are sequence independent).

To design a linear, enthalpy-neutral strand displacement cascade, two parameters are necessary and sufficient: *N* (*N* ⩾ 2) represents the number of double-stranded domains in a fuel; *shift* represents the distance between consecutive single-stranded toeholds (see Figure 1b). *shift* should be between 1 and *N* to ensure signal can be transmitted to the next fuel. Figure 2 shows the diversity of designs when *N* = 6 for all valid values of *shift*.

**Figure 2:**
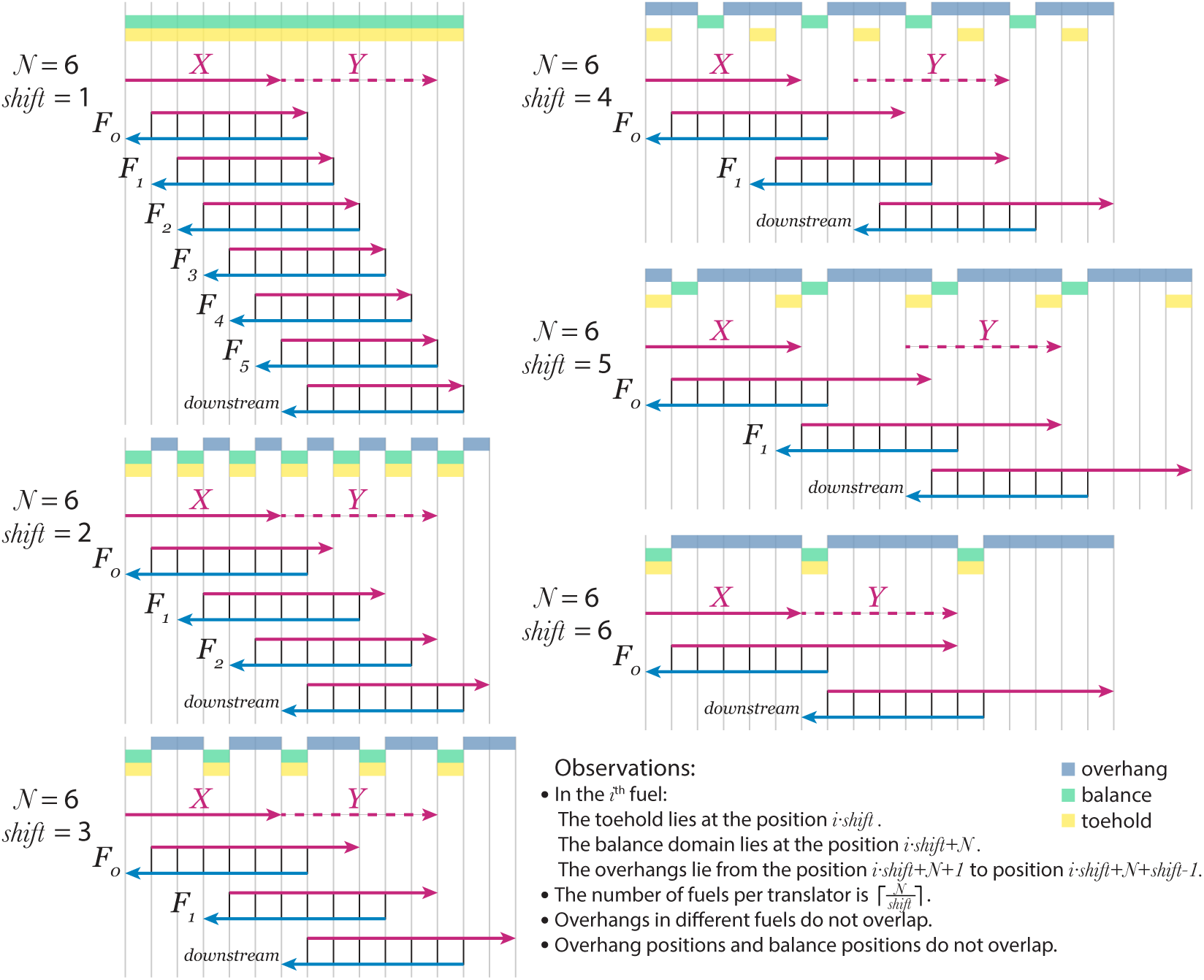
Initial configurations for linear strand displacement cascades with *N* = 6 and all values of *shift*. For each combination of parameters, enough fuels are shown to output a signal of independent sequence from the input *X* (i.e., a full translator), as well as an additional (downstream) fuel. The dashed strand *Y* indicates the positions of the output signal strand that the downstream fuel takes as input. The positions are colored on top of each design according to their position types (see legend). Top strands are colored in purple and bottom strands are colored in blue. Note that although drawn as a single copy, we consider a regime with multiple copies of each fuel. The observations are justified and explained in Section S3.

Using 0 indexing, we assign the domain type starting from the 0^th^ fuel which is responsible for the first strand displacement step (e.g. Figure 1c). The toehold of the 0^th^ fuel is located at position 0. A position can belong to one or more different position types: *toehold, overhang*, and *balance*, depending on which types of domains exist in that position. Each position type has its own color shown in the figures. (We assume the linear strand displacement cascade is extended both upstream and downstream, so that every bottom strand has exactly the same layout of position types on all the domains.) Basic properties and observations for the enthalpy-neutral strand displacement cascade are shown in Figure 2 and listed in Section S3.

### Asymptotic completion level of translator cascades

With a cascade of irreversible reactions, most of the input signal should propagate through to the end. However, enthalpy-neutral cascades are reversible and the amount of output decreases with the length of the cascade since the signal “spreads out” across the layers. Nonetheless, we argue that long cascades are feasible by proving a lower-bound that is *independent of the length of the cascade* on the amount of final signal output.

We simplify each enthalpy-neutral strand displacement reaction to be a bimolecular reversible reaction *X* +*F* ⇌ *Y* +*W*, where *X* is the input signal, *F* is the fuel, *Y* is the displaced strand of this reaction and *W* is the waste species. The enthalpy neutral assumption implies that the equilibrium constant of each reaction can be treated as 1 (Section S4). Now suppose we have a *n*-layer reaction system starting with fuel (*F*_*i*_, *i* = 1, 2, …, *n*) concentrations at 1 and input *X*_1_ at *α*:

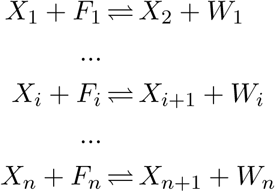

#### Theorem 1.

*Given a linear enthalpy-neutral strand displacement cascades with n (n* ⩾ 2*) layers, and the fuel concentration* 1 *and the input concentration α, the equilibrium concentration of the output signal is at least* 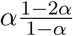 *independent of n*.

The lower bound on the ratio of output to input 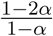 approaches 1 for small *α*; i.e.,almost all of the input is converted to output. For 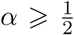 the theorem does not provide apositive lower bound and thus is not meaningful (although larger *α* leads to higher output signal). Since the lower bound is independent of the layer number *n*, concatenating the enthalpy-neutral reactions generates at least a constant fraction of output signal independent of the number of layers.

## Kinetic properties

### Toehold occlusion

Stronger toeholds enable faster initiation of strand displacement [12]; however, toehold strength being too strong slows down reaction kinetics due to undesired toehold binding. Since typically fuels are at high concentration, this is especially problematic when overhangs bind to toeholds of other fuel (when overhang and toehold positions overlap, Figure 3a). This creates so-called *toehold occlusion*, which can significantly slow down the intended reaction [4, 8]. The following theorem characterizes exactly the designs which avoid toehold occlusion.

**Figure 3:**
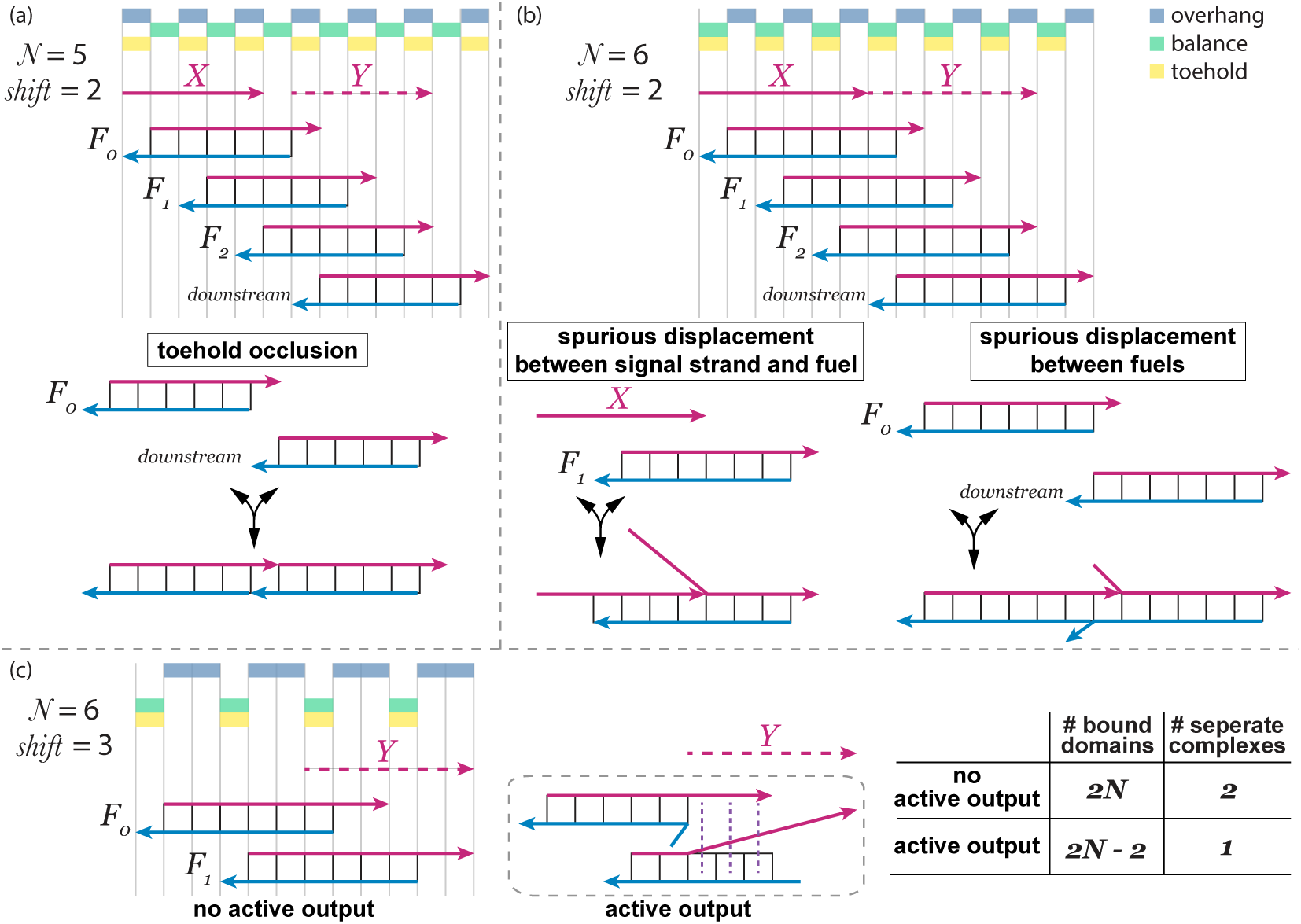
Examples of the properties of linear enthalpy-neutral strand displacement cascades. (a) Toehold occlusion in the design *N* = 5, *shift* = 2. (b) Spurious displacement in the design *N* = 6, *shift* = 2. (c) Energy penalty for active output in the design *N* = 6, *shift* = 3. The dashed gray box indicates the strands in the box are in one complex. The dash purple lines indicate bound domains. The shown active output configuration incurs both an enthalpy (two fewer bound domains) and entropy (one fewer separate complex) penalty and is thus thermodynamically unfavorable.

#### Theorem 2.

*Toehold occlusion is not possible in a linear enthapy-neutral strand displacement cascades if and only if N is a multiple of* shift.

### Spurious strand displacement

Spurious strand displacement can slow down the intended kinetics. Spurious displacement occurs when any proper prefix or suffix of a fuel’ s double-stranded helix is displaced by a spurious invader. A spurious invader of a fuel is any species different from its intended input strand. We analyze spurious displacement between fuels and spurious displacement between a signal strand and a fuel as shown in Figure 3b.

Like toehold occlusion, spurious displacement between fuels can become increasingly problematic at larger concentrations as it makes fuels unavailable for their intended reactions. Figure 4 shows contrasting examples with hard-to-reverse spurious reconfiguration (requiring bimolecular steps to undo) and those in which any spurious reconfiguration can be quickly undone (unimolecular steps are sufficient). To the first approximation, even at the “high” concentrations used here, bimolecular reactions are relatively slower than unimolecular reactions. We prove that in certain cases such hard-to-reverse displacement is thermodynamically penalized (see Section S7). Lemma S5.3 and Lemma S5.4 discuss designs with spurious displacement in the absence of input.

**Figure 4:**
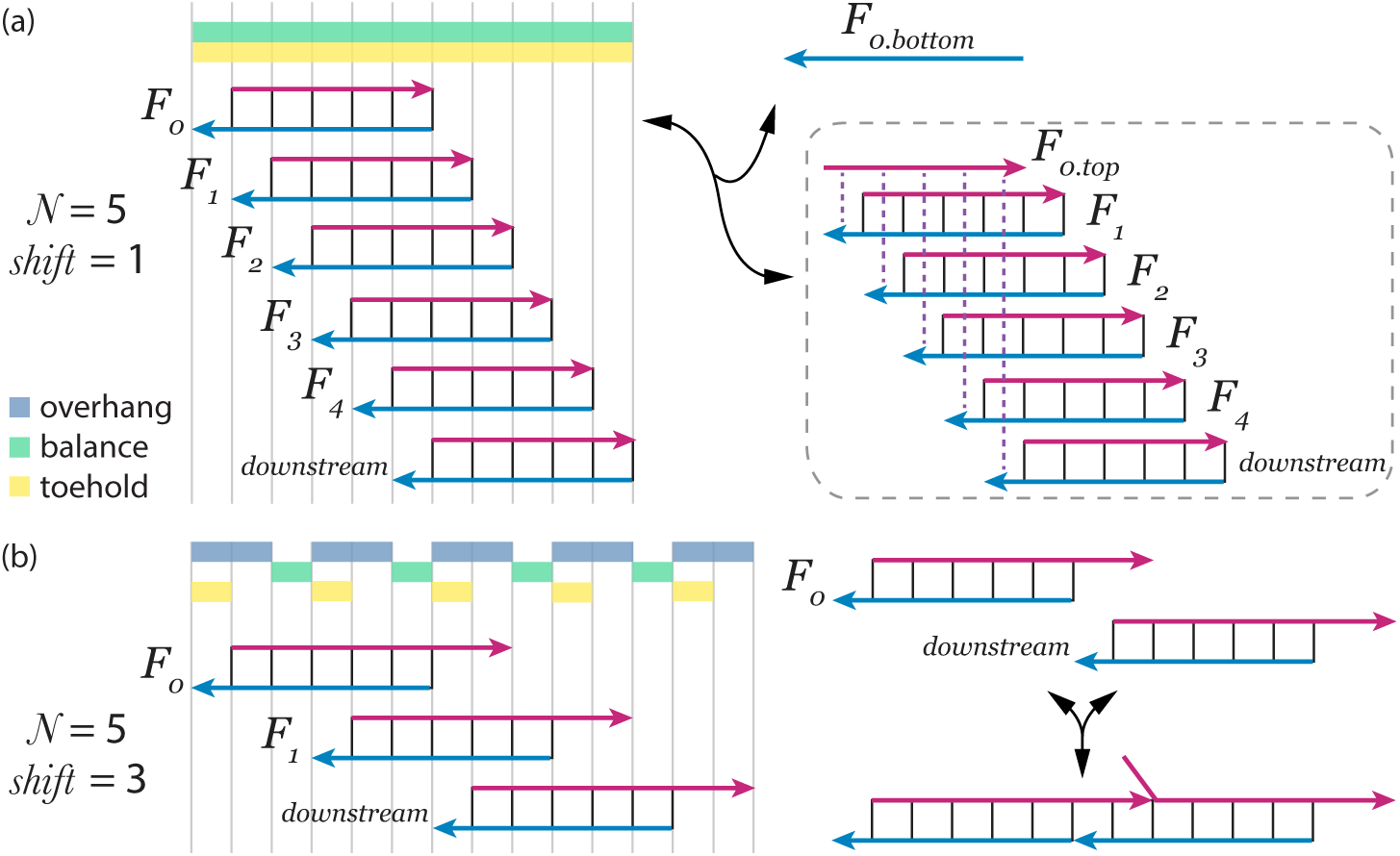
Examples of configurations reachable with spurious strand displacement. (a) Spurious strand displacement results in a configuration that requires a bimolecular reaction to undo. In the absence of input signal, multiple spurious displacement events could result in all of the bottom domains of one fuel being displaced by toeholds of other fuels. This results in a free (unbound) bottom strand. Although the multiple toeless displacement events to cause this reconfiguration are unlikely, once formed, it requires a slow bimolecular reaction to undo. The dashed gray box indicates the strands in the box are in one complex. The dash purple lines indicate bound domains. (b) Spurious strand displacement results in a configuration that requires a unimolecular reaction to undo.

The second type of spurious displacement is when a free signal strand (including the input), can spuriously invade a fuel other than its designed target. When the input concentration is significantly lower than fuel, as is typical, signal strands can become involved in numerous unproductive reactions thus slowing (possibly significantly) signal propagation through every layer of the cascade. Lemma S5.5 discusses spurious displacement in the presence of input.

The following theorem characterizes when both kinds of spurious displacement are avoided.

#### Theorem 3.

*Spurious displacement is not possible in a linear enthapy-neutral strand displacement cascade if and only if* shift = *N* − 1.

### Thermodynamic properties: Thermodynamic penalty to leak

While non-enthalpy neutral strand displacement schemes are typically metastable and leaky if left without input for a long time, we argue that our *EN* design has a programmable *thermodynamic penalty* to leak (comparison details in Section S8).

Our thermodynamic arguments are based on “enthalpy”—counting the number of bonds formed (more bonds are more favorable) and “entropy”—the number of separate complexes (more complexes are more favorable) [21, 22, 23]. We acknowledge that our use of “enthalpy” and “entropy” are merely shorthand; there are entropy contributions of bond formation, and there are other components of entropy such as conformational entropy and the distinguishability of molecules.

An example of analyzing the energy penalty to leak is shown in Figure 3c. Although strand displacement systems are designed to not generate output assuming displacement only follows toehold binding, leak can result from “toeless” strand displacement reactions caused by spontaneous fraying at the ends of helices [24]. Note that the newly exposed single-stranded region (corresponding to *Y*) in the figure can subsequently undergo strand displacement with downstream fuels. Varying design parameters *N* and *shift* changes the thermodynamic penalty (according to the two thermodynamic factors) to leak. All of our combinatorial arguments are valid both in the single-molecule regime or when the fuels have multiple copies. Different fuels may also be present in different amounts.

For the purposes of the arguments, we say the *ith-output region* (for any *i >* 0) consists of *N* positions *i*·*shift* to *i*·*shift*+*N* −1. A *configuration* is a matching between corresponding top and bottom domains, where each match is one bond. A configuration has an *active output* if some complex has open top domains in all *N* positions of some output region. These positions represent the domains in a top strand that can displace a strand of the downstream fuel. In the *initial configuration* each fuel complex is separate and unreacted (e.g. the configurations shown in Figure 2).

We first show that in the absence of input, the initial configurations are the thermodynamically preferred configurations for designs without toehold occlusion. Thus the thermodynamic penalty to produce an active output in the absence of input can be properly based on the comparison between the active-output configuration and the initial configuration.

### Initial configuration is thermodynamically preferred without toehold occlusion

There might be tradeoffs between enthalpy (number of bonds) and entropy (number of seprate complexes), but if we find a configuration that simultaneously maximizes both, it is thermodynamically preferred. In our case, we will show that the initial configuration simultaneously maximizes both unless we are willing to lose a lot (*N*) of bonds. Thus it is the thermodynamically preferred configuration in a wide range of experimental regimes.

In our proof of the following theorem, we gradually restrict the possible complexes of the thermodynamically preferred configuration through Lemma S6.2, Lemma S6.3, and finally by showing that they must consist of exactly one top and one bottom strand (Lemma S6.4). Combined with an analysis of the tradeoff between bonds and separate complexes later developed in Theorem 6, this yields the following result:

#### Theorem 4.

*For designs without toehold occlusion, given a linear enthalpy-neutral strand displacement cascade of arbitrary depth, in the absence of input, we have: (i) Among the configurations with maximum bonding, the initial configuration is the unique configuration that maximizes the number of separate complexes. (ii) The initial configuration has at least N more bonds than any configuration with more separate complexes*.

### Enthalpic penalty to leak

Our arguments about the enthalpic penalty are based solely on counting the difference in bonds between the thermodynamically preferred configuration and the configuration with active output. We show that the enthalpic penalty to leak can be increased arbitrarily by tuning the parameters *N* and *shift*.

To have an active output, it is necessary to break some bonds if there is no free top domain among the output region. Thus it is sufficient to count the number of positions among the output region that do not have excess top domains (top-limiting positions; Lemma S6.6).

#### Theorem 5.

*Given a linear enthalpy-neutral strand displacement cascade of arbitrary depth, in the absence of input, any configuration having active output has at least* 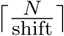*fewer bonds than the maximum-bond configuration*.

This theorem implies that in the absence of input there is an enthalpic penalty of 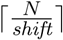 bonds to generate an active output from a maximum-bond configuration. Note that in the designs without toehold occlusion, the initial configuration maximizes the number of bonds and so can act as the unique point of comparison (via Theorem 4). The theorem still applies to designs with toehold occlusion but the thermodynamically preferred configurations are not necessarily maximum-bonded so the relevance of the point of comparison is not clear.

In contrast, when the input is present, the signal can be propagated all the way with no net enthalpic cost. Thus by increasing *N* we can enlarge the enthalpic penalty to leak without increasing the enthalpic penalty to correct output. The enthalpic penalty holds even at high concentrations when there is little entropic penalty of joining two complexes into one, and thus sets the lower bound on the required energy for leak independent of concentration.

The enthalpic penalty shows that leak results in fewer bonds compared with maximum bonding. Moreover, Theorem 4(ii) shows that leak cannot compensate for the loss of bonds with the thermodynamic benefit of creating additional separate complexes (unless *N* bonds break). Indeed, in the next section we show that leak results in a *decrease* in the number of separate complexes, and that this entropic penalty can be made arbitrarily large by varying *N* and shift.

### Entropic penalty to leak

In this section we show that no matter how strands are rearranged in our enthalpy-neutral strand displacement cascades, any configuration with active output has fewer separate complexes— thus it incurs an entropy penalty. As the point of comparison, we count the difference in the number of separate complexes between the thermodynamically preferred configuration as captured by Theorem 4 and the active-output configuration.

By Theorem 5, we know that the enthalpic penalty has a lower bound of 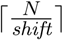. Thus the arguments below assume that the configuration with active output has *r* bonds fewer than maximum, where 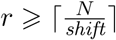. Beyond this lower-bound on *r*, the entropy penalty can be traded-off for an decreased enthalpy penalty by increasing *r*. Intuitively, forcing fewer bonds to form gives more flexibility to produce active output without bringing as many complexes together.

To count the difference in the number of separate complexes, we can equivalently count the number of top strands associated in a complex in the active-output configuration (since the thermodynamically preferred configuration has only one top strand in every complex by Lemma S6.4). Then by a combinatorial argument based on counting the types of positions in top and bottom strands (Lemma S6.9, Lemma S6.10, Lemma S6.11 and Lemma S6.12), we show that for active output there must exist some complex with at least 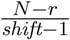top strands (Lemma S6.11 and Lemma S6.12), implying the following theorem:

#### Theorem 6.

*For designs with* shift ≠ 1 *and no toehold occlusion, given a linear enthalpy-neutral strand displacement cascade of arbitrary depth, in the absence of input, any configuration with active output and r (r* ∈ N, *r* ⩾ 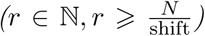*) bonds away from maximum has at least* 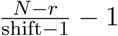 *fewer separate complexes than the initial configuration*.

This theorem implies that in the absence of input there is an entropic penalty of 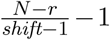 units to produce an active output. In the case of *shift* = 1, active output requires an enthalpy penalty of *r* = *N*, but can result in one additional separate complex.

As in the case of enthalpy, when the input is present, the signal can be propagated all the way with no net entropic cost. Combined with the results of the previous sections, if we fix *shift* and increase *N*, both the enthalpic (Theorem 5) and entropic penalty (Theorem 6) to leak increases, while the desired triggering signal reaches an asymptote (Theorem 1).

### Trade-offs between the properties

We have shown that different choices of *N* and *shift* yield *EN* designs with varying thermodynamic and kinetic properties. We desire a large thermodynamic penalty to leak, which requires a large ratio *N/shift* (Theorems 5 and 6). However, only designs with *shift* = *N* − 1 can avoid spurious strand displacement 3, which is incompatible with a large *N/shift* ratio. In fact, designs with the largest thermodynamic penalty to leak also have the most potential spurious interactions between fuels and signal strands. (Compare the spurious interactions of the *X* input signal with fuels other than its intended target for different *shift* in Figure 2.)

Besides spurious displacement, the other undesired kinetic property is toehold occlusion. While the absence of toehold occlusion is compatible with a large thermodynamic penalty (Theorem 2), there is a fundamental incompatibility between the two kinetic properties and only one design can avoid both:

#### Corollary 1.

*The only design that avoids both toehold occlusion and spurious displacement is N* = 2, shift = 1.

In summary, there is no best design with respect to all thermodynamic and kinetic properties studied here. Instead, the entire taxonomy we develop informs the choice of translator design, and one should be chosen based on the expected conditions of its planned use. For example, high concentration conditions may be best served by a design with no toehold occlusion and a balance between its enthalpic penalty to leak and its potential number of spurious displacement reactions.

## Experiment results

### Designs with different energy penalties to leak

We experimentally tested the energy penalties to leak among two sets of design parameters (Figure 5). The *N* = 6, *shift* = 3 design has 2 unit of enthalpic cost and 1 unit of entropic cost to leak. The *N* = 6, *shift* = 6 design has only 1 unit of enthalpic cost, and no entropic cost. We used fluorophore and quencher labeled “reporter” complex to detect output signal. Experiments confirm that the *N* = 6, *shift* = 3 design has less leak than the *N* = 6, *shift* = 6 design (Figure 5). The leak concentration of the *N* = 6, *shift* = 6 design increases with temperature as is consistent with an enthalpic penalty. In contrast, the effect of entropy on the equilibrium should be independent of temperature. Leak in the *N* = 6, *shift* = 3 design does not have significant temperature dependence, which may be due to the entropic contribution to the leak penalty.

**Figure 5:**
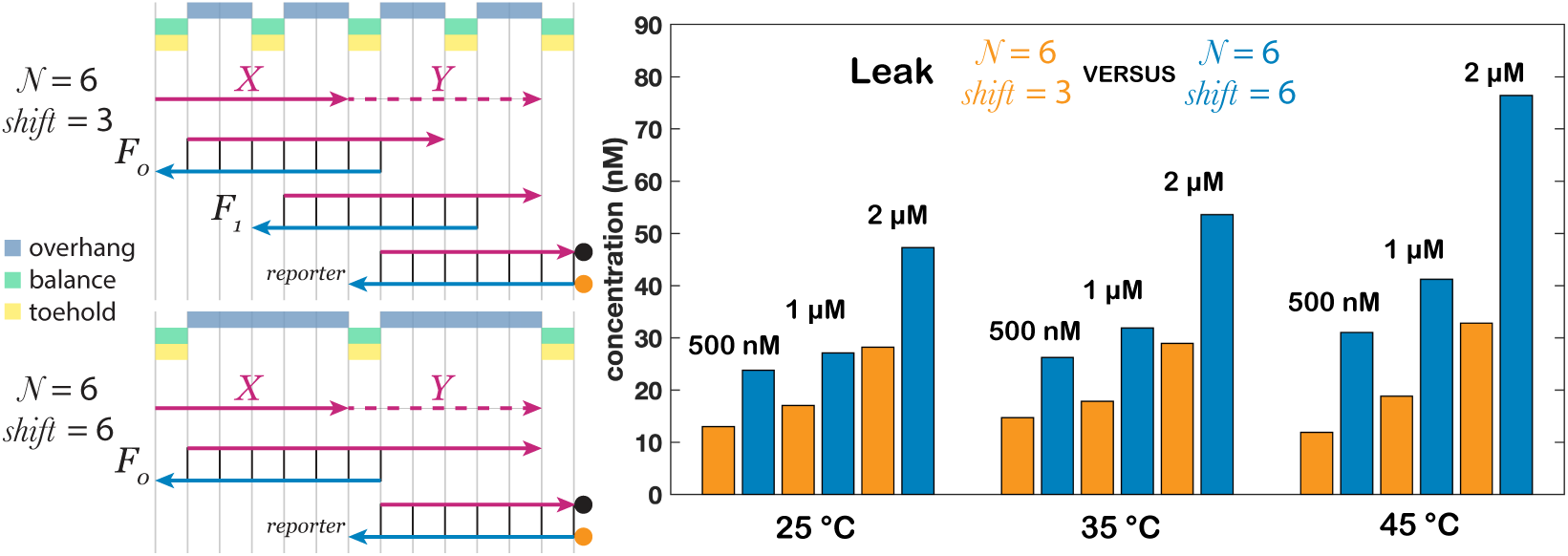
Comparison of designs with different energy penalties to leak. Signals at different thermodynamic equilibrium (25 °C, 35 °C and 45 °C) after a slow annealing from 90 °C (0.06 °C/min) without input strand. The design with parameter set *N* = 6, *shift* = 3 has less relative leak. The top strand of the reporter is labeled with a quencher (black circle). The bottom strand of the reporter is labeled with a fluorophore (orange circle). The concentrations for fuels and reporters are labeled in the figure.

Since the energy penalty to leak of the *N* = 6, *shift* = 6 design is not entropic, the leak concentration is expected to be proportional to the initial concentration of the reacting species. However, the experimental leak concentration does not double when the initial concentration doubles. This deviation could be caused by some reactions changing entropy that are not captured in our model, such as sequence-dependent spurious binding or the fluorophore-quencher interaction. (Note that this design is predicted to not have toehold occlusion.)

In the presence of input signal, the two designs produce similar amounts of output signals (Figure S3). Indeed, the low-leak *N* = 6, *shift* = 3 design has slightly higher output.

### Designs with and without toehold occlusion

Our theoretical results classify designs with toehold occlusion (Theorem 2), which can hinder the kinetics of desired strand displacement. Here, we experimentally compare the designs with (*N* = 5, *shift* = 2) and without (*N* = 6, *shift* = 2) toehold occlusion. As shown in Figure 6, at various concentrations of input, the design without toehold occlusion (*N* = 6, *shift* = 2) reaches completion faster than the design with toehold occlusion (*N* = 5, *shift* = 2). When the input concentration is 1 *µ*M, the design without toehold occlusion reaches 86% of completion before the first measured data point, but it takes nine minutes for the design with toehold occlusion to reach the same completion level. Note that these two designs share the same toeholds, but there are slight differences between the energies of the toehold and balance domains. These thermodynamic differences actually favor the design with toehold occlusion, and thus cannot account for the kinetic slowdown.

**Figure 6:**
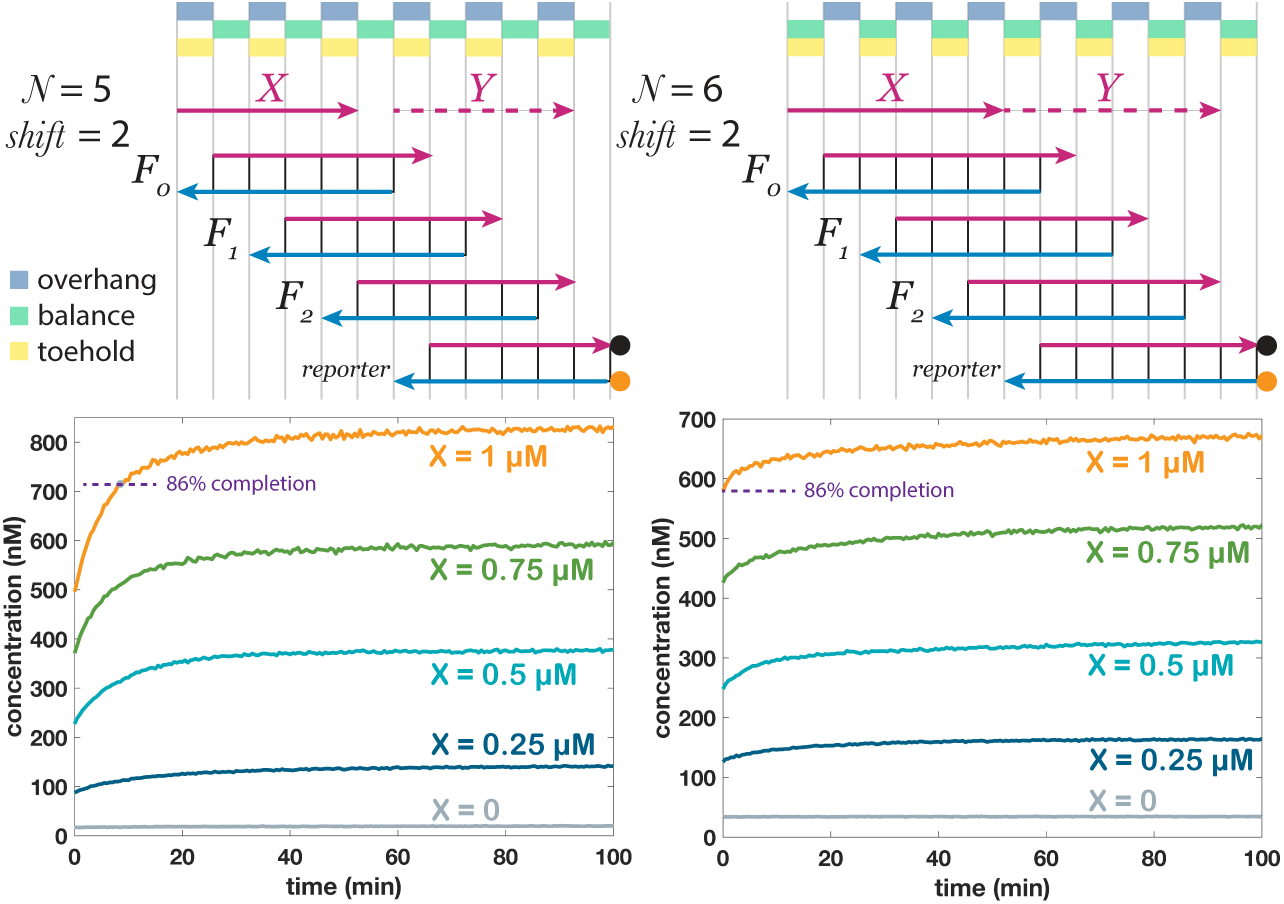
Comparison of kinetics of designs with (*N* = 5, *shift* = 2) and without (*N* = 6, *shift* = 2) toehold occlusion. We define completion as the signal level at 100 min. The design with the parameter set *N* = 6, *shift* = 2 is significantly faster. The concentrations of reporter and fuels were 2 *µ*M. The reaction temperature was 25 °C. As in all kinetic curves in this work, time 0 represents the first data point measured, which is about 2 minutes after samples were mixed.

### Spurious displacement pathway with signal strands

Spurious displacement may result in the formation of complex multi-stranded structures or intermediates, which could interfere with desired kinetics. Theorem 3 classifies designs with and without spurious displacement overall, and Lemmas S5.4 and S5.5 separate the cases of spurious displacement with the input absent or present. We chose the design with *N* = 6, *shift* = 1, which falls into the category of spurious displacement with input, to test the predicted spurious displacement reaction pathway. This parameter set represents the extreme case of spurious displacement since every input to a fuel has multiple invading sites with other fuels.

Figure 7 shows two possible spurious displacement pathways involved with signal strands and the fuels that are not the designed targets of the signal strands. The signal strand first binds and partially displaces the top strand of the fuel, exposing a single-stranded portion of the top strand. The single-stranded portion can then interact with other downstream species. We use a reporter to represent the downstream species and detect the reaction pathway. After a 3-way branch migration and then a 4-way branch migration process, the fluorophore and quencher in the reporter separate. The gradual increase of fluorescence only in the presence of both an upstream fuel and a signal that can spuriously react with the fuel is consistent with the described spurious displacement pathway.

**Figure 7:**
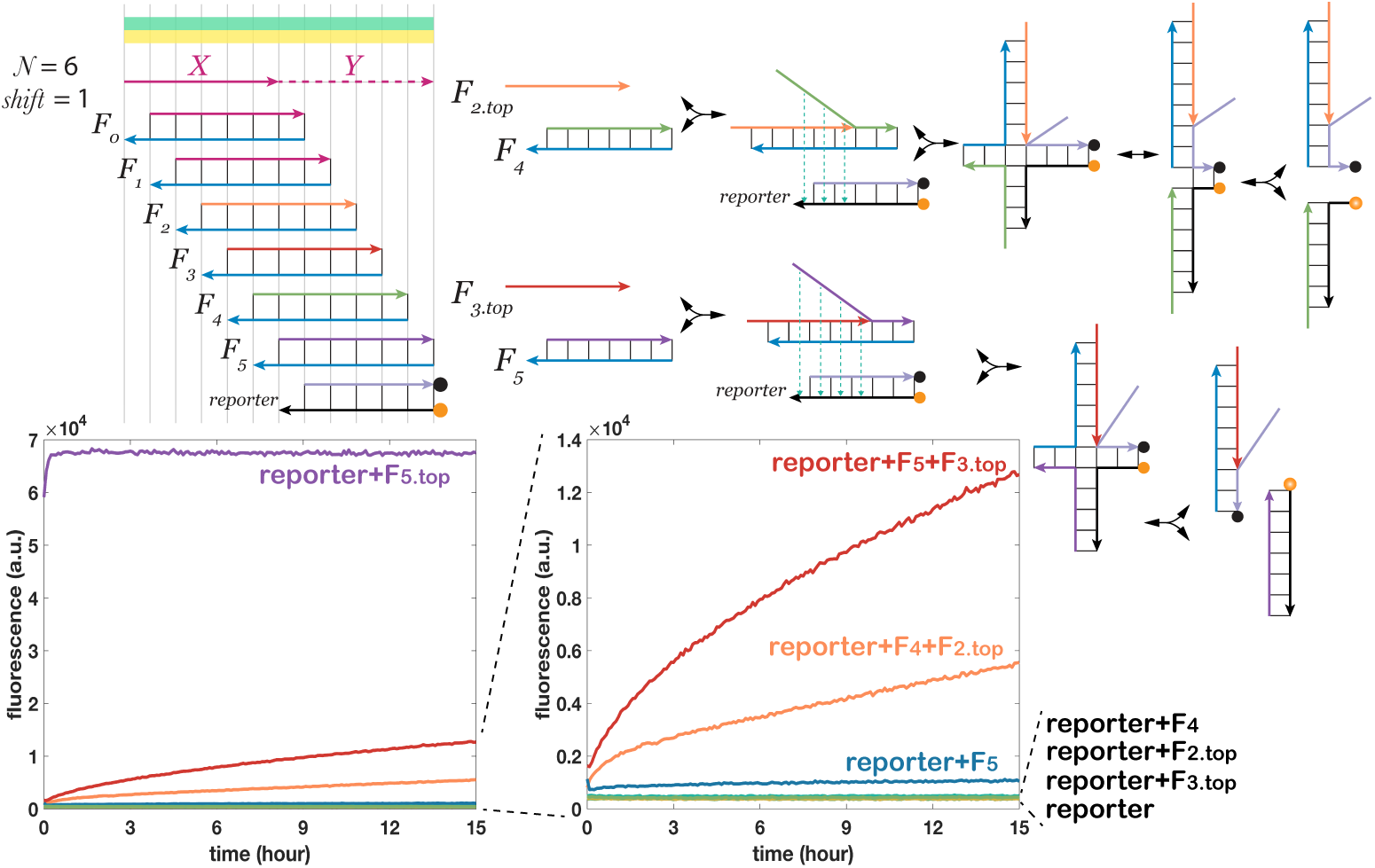
Spurious displacement pathways and kinetics of spurious interactions at the design *N* = 6, *shift* = 1. The concentrations of reporter and fuels were 500 nM. The input at each layer was 100 nM. The reaction temperature was 25 °C.

### Leak reduction compared to designs with only an entropy barrier

Prior work (*NLD* design) based leak reduction on a kinetic barrier wherein multiple separate complexes needed to co-localize to generate leak (entropy barrier) [9, 10]. Our *EN* design has a stronger guarantee of correct behavior in two ways: (1) by substituting an overall thermodynamic penalty for a kinetic barrier, (2) by basing this penalty on both entropy and enthalpy. To experimentally compare leak reduction of our *EN* design to the prior work, we chose one of the parameters *N* = 5 and *shift* = 3 with the desired property that leak requires at least two units of enthalpic penalty (breaking two bonds) compared with the maximum-bond configuration (Theorem 5). We compare with the typical leaky translator with no leak reduction (*SLD* design) and the leak reducing *NLD* design with *N* = 2 (*DLD* design) [9, 10] (Figure 8a).

**Figure 8:**
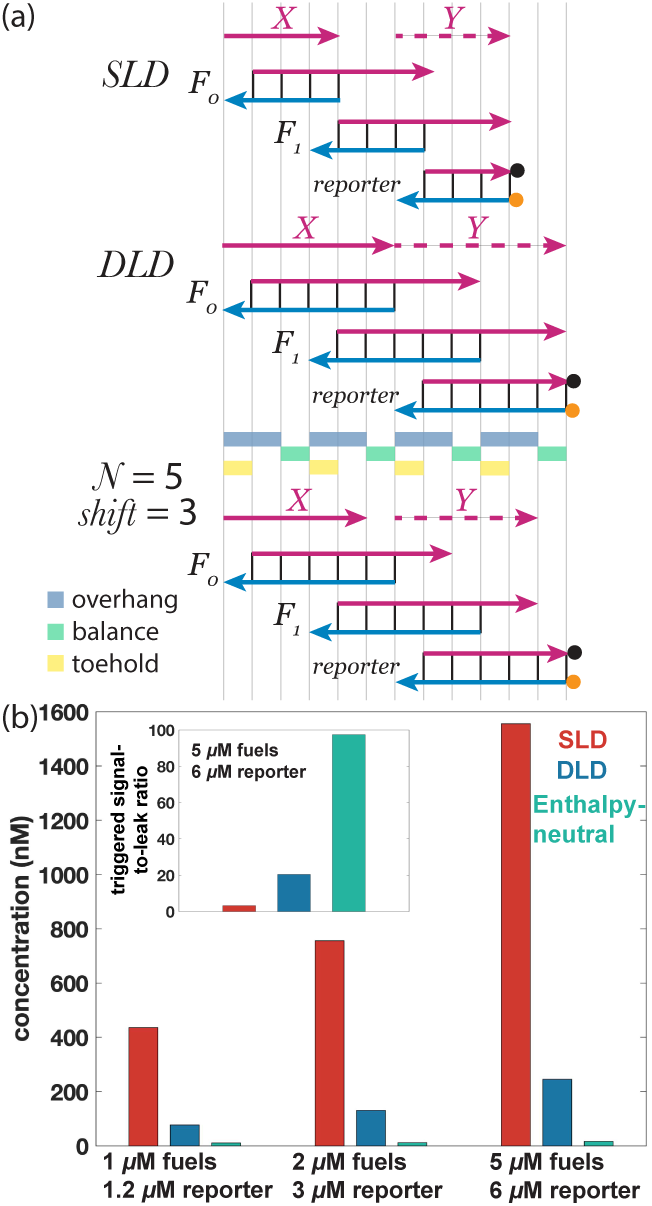
Comparison of leak between (a) the *SLD, DLD* and *EN* design (*N* = 5, *shift* = 3) from the perspective of (b) the thermodynamic equilibrium. The experiments measuring the leak concentration at the equilibrium do not contain the input signal strands. Fuels and reporter were slowly annealed (0.06 °C/min) from 90 °C to 25 °C within 18 h. (Inset) The triggered signal-to-leak ratios of the *NLD* design and the *EN* design. The *SLD* and the *DLD* designs were assumed to reach full completion. The triggered signal for the *EN* design was shown in Figure S4. The concentrations for fuels and input were 5 *µ*M, and the reporter 6 *µ*M. The reaction temperature was 25 °C. For simplicity, the clamp domains in the *SLD* and *DLD* designs (which extend the bottom strand by 2 nt) are not shown here but are illustrated in Figure S8.

For a most fair comparison, in our experiments, these three translator designs share the same sequence space and all the fuels have the same toeholds. Moreover, the *DLD* and the *EN* design share the same reporter complex. The detailed comparison for the designs at the sequence level is shown in Figure S8. Figure 8b compares the amount of leak at thermodynamic equilibrium after annealing. The *EN* design was observed to have an order of magnitude less leak than the *DLD* design and about two orders of magnitude less than the *SLD* design. The fuel concentrations used here are 10–100 times larger than the typical concentrations used in strand displacement systems, showing that *EN* schemes can reduce leak in the fast (high-concentration) regimes.

Although enthalpy-neutral strand displacement is not driven by the formation of new bonds as are the *SLD* and *DLD* designs, we observed that the triggered signal is only around three times lower than full triggering. Thus the signal-to-leak ratio of the *EN* design is 30 times higher than of the *SLD* design and four times higher than of the *DLD* design (Figure 8b inset).

### Desired triggering and leak in longer cascades

Beyond a single translator, we wanted to know (1) how much leak (without input signal) and (2) desired output signal (with input signal) a translator cascade can generate as the number of translators increases.

Figure 9a shows that in the time period of the experiment, in the absence of input signal strand, the translator cascades of 1 to 6 fuels (1 to 3 translators) all show no apparent leak. In the presence of input signal, the completion level decreases with the number of layers. However, as more layers are added, the completion level does not decrease linearly, and indeed seems to approach an asymptote—behavior consistent with the theoretical prediction in the previous section.

**Figure 9:**
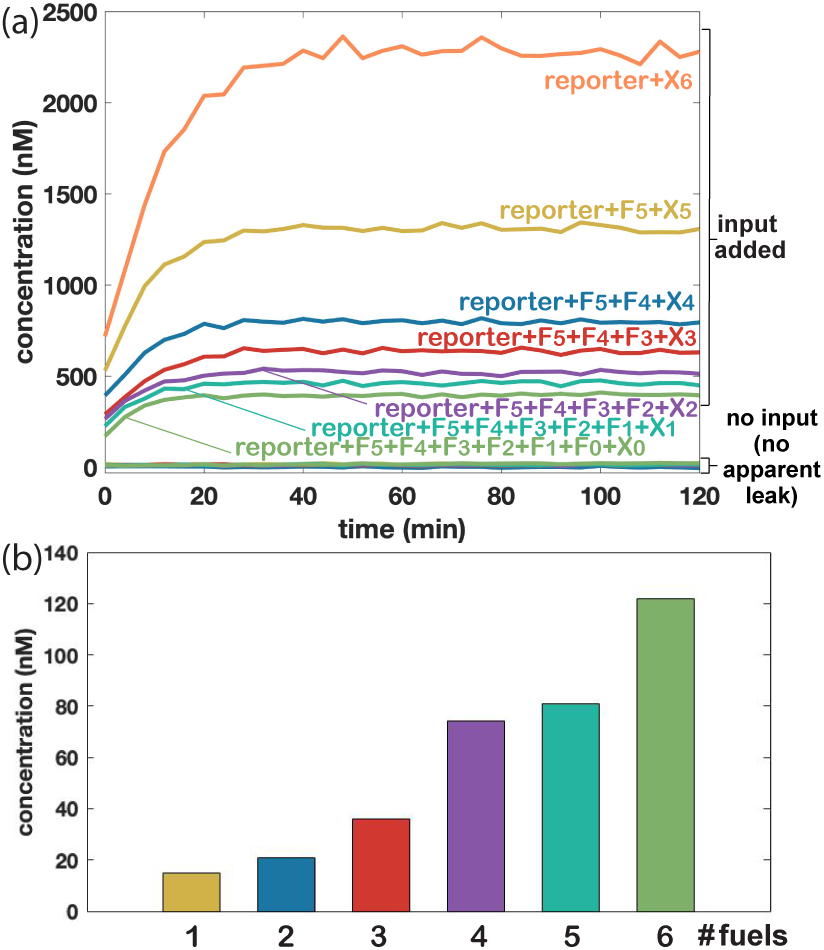
Kinetics and thermodynamic equilibrium of translator cascades with the *EN* design (*N* = 5, *shift* = 3). (a) Kinetic behaviour in the presence and absence of input signal, for cascades of different length. (b) The total amount of leak in the absence of input signal at thermodynamic equilibrium measured after annealing, for cascades of different length. The concentrations of the reporter and the fuels were around 5 *µ*M. The concentration of each input was 2.5 *µ*M. The reaction temperature was 37 °C. Note that *X*_*i*_ in the figure refers to the input to fuel *F*_*i*_, and *X*_6_ is the input to the reporter. (Note that the actual reporter used is one domain shorter than the *N* = 5 design and the overhand on *F*_5_ was shortened to compensate (Figure S9), which is not expected to affect the completion level.)

Figure 9b studies the leak of translator cascades of varying depth at thermodynamic equilibrium. The leak at equilibrium increases as the number of fuels increases. However, even if there are 6 fuels (3 translators), the total leak concentration is still less than 3% of the fuel concentration. Indeed, it is about 5 times less than the leak concentration for a *single* translator in the *DLD* design, and it is an order of magnitude less than that in the *SLD* design. Note that although the absolute amount of triggered signal of the *EN* design is lower than that of the *DLD* design, the relative triggered signal-to-leak ratio of the *EN* design is still higher than the *DLD* design. The results suggest that the *EN* design could be preferable, especially when absolutely smaller leak is required, such as when concatenating the translators with downstream catalytic or auto-catalytic systems.

## Discussion

The design space of enthalpy-neutral strand displacement cascades shows a surprising range in terms of kinetic and thermodynamic behaviors which was previously not explored. The full diversity of enthalpy-neutral linear-topology strand displacement cascades is accessible by varying two parameters *N* and *shift*, with different susceptibilities to toehold occlusion and spurious displacement, and rigorous guarantees on the enthalpic and entropic penalties to leak determined by these parameters. We formalized the analysis describing how the enthalpic and entropic penalties to leak can be raised arbitrarily, captured the inherent tradeoff between the two penalties. The enthalpic penalty argument is particularly germane for the high concentration regime where the entropy penalty for joining complexes is smaller. We further proved that certain parameter values result in other desirable properties like no spurious displacement and no toehold occlusion. Since no design satisfies all desired properties, understanding the taxonomy of designs is necessarily to tune the tradeoffs and make proper design choices.

In addition to theoretical analysis (Section S8), we experimentally tested designs with different energy penalties to leak, confirming different degrees of leak reduction and improvement over the previous *NLD* design. Our experiments also corroborated that slower kinetics due to toehold occlusion in otherwise similar designs. Finally, in identifying spurious displacement reaction pathways, the experiments confirm the rationale for studying spurious displacement.

The scale of current strand displacement systems exceeds the capabilities of thermodynamic modeling tools [25] and (even more so) of kinetic simulators [26]. Thus it is challenging to understand large-scale molecular behaviors due to the increasing complexity. However, combinatorial arguments like those espoused in this work apply to infinitely many, arbitrarily large systems. This kind of analysis is analogous to rigorous proofs of algorithm correctness in computer science, in which a single proof applies to all possible inputs that the algorithm may have, and such lines of reasoning could be developed for broader classes of engineered molecular systems.

## Methods

### Proofs

All the proofs are in SI.

### Sequence design

The details for DNA sequence design principles are in SI.

### DNA oligonucleotides

DNA oligos were synthesized by Integrated DNA Technologies (IDT). The unlabelled oligonucleotides were purchased PAGE purified by IDT. The oligonucleotides with fluorophore or quencher modifications were ordered HPLC (high-performance liquid chromatography) purified by IDT. Upon arrival, these DNA oligonucleotides were suspended in Milli-Q water. The concentration of each strand was quantified by NanoDrop. The absorbance at 260 nm was recorded and the concentration (*c*) was calculated as *c* =[Absorbance]*/e*, where *e* is the extinction coefficient provided by IDT.

### Fuel and reporter preparation

Fuels and reporters were prepared according to reference [10]. Fuels and reporters were annealed in TAE/Mg^2+^ buffer (40 mM Tris, 20 mM acetic acid, 1 mM EDTA, and 12.5 mM Mg^2+^, pH ≈ 8.0) with 10% excess of top strands from 90 °C to 25 °C within 1.5 *h*. PAGE purification was then conducted to remove malformed structures or single-stranded DNA [4] from fuels. Purified duplexes were quantified by NanoDrop again and also concentrated by centrifugal filters (Amicon Ultra-0.5 mL, 10K device) to achieve a high stock concentration. Reporters were not purified. The extinction coefficients for complexes were estimated the same way as in reference [10]: adding up the estimated extinction coefficients for the single-stranded and double-stranded parts.

### Fluorescence measurement

Experiments for the enthalpy-neutral DNA strand displacement cascades with parameters *N* = 5, *shift* = 3 were measured by BioTek Cytation 5. All other experiments were measured by BioTek Synergy H1. The NBS (non-binding surface) 384 well plates with clear flat bottom (Corning # 3544) were used. Fluorescence was measured from the bottom. The excitation/emission wavelengths were set to 575/610 nm for the fluorophore ROX. For Cytation 5 measurement, the excitation bandwidth was fixed at 9 nm and the emission bandwidth was fixed at 20 nm.

For kinetic experiments, all the samples were mixed and prepared at room temperature (close to 25 °C), and then quickly transferred to a plate (within 2 minutes). Time 0 in plots represents the moment the first data point was measured.

For equilibrium experiments, all the samples were annealed from 90 °C to desired temperatures with 20 minutes per degree. The plate was incubated in the plate reader at the desired temperatures for 10 minutes prior to data collection to let the samples reach the same temperature as the experimental setting.

## Data fitting and normalization

The details of data fitting and normalization are in SI.

## Acknowledgement

The authors thank Cameron Chalk for reading and commenting on the manuscript. B.W. and D.S. were supported by NSF grants CCF-1618895 and CCF-1652824. C.T. was supported by NSF grants CCF-1317694 and CCF-2106695.

## Author Contributions

B.W. initiated the project; B.W., C.T. and D.S. designed the research; C.T. proved the theorems for kinetic properties; B.W. and D.S. proved the other theorems; B.W. performed experiments and analyzed data; B.W., C.T., D.S. wrote the paper.

