## Supporting Information for "Speed and correctness guarantees for programmable enthalpy-neutral DNA reactions"

#### Contents

**S1 One unit of enthalpic/entropic penalty** **3**

**S2 Useful Lemmas** **3**

**S3 Observations and properties for linear enthalpy-neutral strand displacement cascade** **4**

**S4 Proof for Asymptotic completion level of translator cascades** **5**

**S5 Proofs for kinetic properties** **6**

**S9 Orthogonal toeholds** **18**

**S10 Sequence design** **19**

**S11 Data fitting and normalization** **20**

**S12 Triggered signals for designs with different energy penalty to leak** **22**

**S13 Comparison of triggered signals between the prior *NLD* designs and the enthalpy-neutral strand displacement design** **23**

**S14 Sequences** **24**

#### S1 One unit of enthalpic/entropic penalty

Roughly speaking, "one unit of enthalpic penalty" corresponds to an average of $l \cdot 1.5$ kcal/mol, where $l$ is the length of the domain (typically 5-10 nucleotides for a toehold). "One unit of entropic penalty" at concentration $C$ M corresponds to $\Delta G^{\circ}_{\text{assoc}} + RT\ln(1/C) \approx 1.96 + 0.6\ln(1/C)$ kcal/mol [1]. With these numbers, at roughly 650 nM concentration, binding an additional $l = 7$ domain is equal to one unit of entropy. At low concentrations the entropic penalty becomes dominant, while the enthalpic penalty prevails at high concentrations.

#### S2 Useful Lemmas

We prove properties of designs based on their parametrization of $N$ and *shift*. Many of our arguments rely on showing whether regularly spaced positions with certain domains (such as toeholds) can intersect with other regularly spaced positions or intervals with different domains (such as overhangs). To simplify those arguments we first establish the following claims.

**Lemma S2.1.** *Let $p$, $q$, and $r$ be natural numbers with $p > 0$ and $r - q > 0$. $\forall i \in \mathbb{N}, \exists j \in \mathbb{N}$ such that $j \cdot p + q = i \cdot p + r$ if and only if $r - q$ is a multiple of $p$.*

*Proof.* Fix any $i \in \mathbb{N}$. Suppose $j \cdot p + q = i \cdot p + r$, for some $j \in \mathbb{N}$. Then $j = i + \frac{r-q}{p}$ and $r - q$ must be a multiple of $p$. □

**Lemma S2.2.** *Let $p$, $q$, and $r$ be natural numbers with $p > 1$ and $r - q > 0$. $\forall i \in \mathbb{N}, \exists j \in \mathbb{N}$ such that $j \cdot p + q$ is contained in the interval $[i \cdot p + r, (i+1) \cdot p + r - 2]$ if and only if $r - q - 1 > 0$ and $r - q - 1$ is not a multiple of $p$.*

*Proof.* Call $k \cdot p + q$ a *valid position*, for any $k \in \mathbb{N}$. Fix any $i \in \mathbb{N}$. First we consider the case when $r - q - 1 = 0$ (and thus $r = q + 1$). Consider any interval $[i \cdot p + r, (i+1) \cdot p + r - 2]$=$[i \cdot p + q + 1, (i+1) \cdot p + q - 1]$. Suppose by contradiction that a valid position $j \cdot p + q$ intersects the interval. Then we have the following: (i) $i \cdot p + q + 1 \leq j \cdot p + q$ which simplifies to $j \geq i + \frac{1}{p}$ and thus $j \geq i + 1$ (since $j \in \mathbb{N}$, and $p > 0$). (ii) $j \cdot p + q \leq (i+1) \cdot p + q - 1$ which simplifies to $j \leq i + 1 - \frac{1}{p} < i + 1$. Contradiction.

Finally, consider two cases when $(r - q - 1) > 0$: (1) $(r - q - 1)$ is a multiple of $p$. Then $i \cdot p + q + (r - q - 1) = i \cdot p + r - 1$ is a valid position and the next valid position occurs at $(i+1) \cdot p + r - 1$. Thus there is no valid position in the interval $[i \cdot p + r, (i+1) \cdot p + r - 2]$. (2) $(r - q - 1)$ is not a multiple of $p$. Let $\delta$ be the remainder; $1 \leq \delta \leq p - 1$. Thus the smallest valid position larger than $i \cdot p + q + (r - q - 1)$ occurs at $i \cdot p + r - 1 + \delta$. Since $1 \leq \delta \leq p - 1$, we know that it falls in the interval $[i \cdot p + r, (i+1) \cdot p + r - 2]$. □

### S3 Observations and properties for linear enthalpy-neutral strand displacement cascade

Since every desired reaction is a toehold exchange strand displacement reaction (i.e., there is a balance domain), we have the following observations and properties:

**Observation S3.1.** *The number of domains comprising an overhang in a fuel is* $\text{shift} - 1$.

The positions of the domain types are listed below.

**Observation S3.2.** *In the $i^{th}$ fuel, (a) the toehold lies at the position $i \cdot \text{shift}$; (b) the balance domain lies at the position $i \cdot \text{shift} + N$; (c) the overhangs lie from the position $i \cdot \text{shift} + N + 1$ to position $i \cdot \text{shift} + N + \text{shift} - 1$.*

The number of fuels per translator is $\lfloor \frac{N+\mathit{overhang}}{\mathit{shift}} \rfloor = \lfloor \frac{N+\mathit{shift}-1}{\mathit{shift}} \rfloor = \lfloor \frac{N-1}{\mathit{shift}} \rfloor + 1 = \lceil \frac{N}{\mathit{shift}} \rceil$, obtaining the following observation:

**Observation S3.3.** *To get an output that is sequence independent of the input, the number of fuels needed is* $\lceil \frac{N}{\text{shift}} \rceil$.

**Lemma S3.4.** *Overhangs in different fuels do not overlap.*

*Proof.* By Observation S3.2, the overhang positions in fuel $i$ are between $i \cdot \mathit{shift} + N + 1$ to $i \cdot \mathit{shift} + N + \mathit{shift} - 1$. So the $i^{th}$ fuel's rightmost overhang position ($i \cdot \mathit{shift} + N + \mathit{shift} - 1$) is smaller than the $i + 1^{th}$ fuel's leftmost overhang position ($i \cdot \mathit{shift} + N + \mathit{shift} + 1$). □

**Lemma S3.5.** *Overhang positions and balance positions do not overlap.*

*Proof.* If $\mathit{shift} = 1$ there are no overhangs and we are done. Suppose $\mathit{shift} > 1$. The $i^{\text{th}}$ balance domain lies at position $i \cdot \mathit{shift} + N$. The $j^{\text{th}}$ overhang lies at positions between $j \cdot \mathit{shift} + N + 1$ and $(j + 1) \cdot \mathit{shift} + N - 1$. Let $r = N + 1$, $p = \mathit{shift}$ and $q = N$. By Lemma S2.2, a position that has a balance domain cannot intersect a position that has an overhang since $r - q - 1 = 0$. □

#### S4 Proof for Asymptotic completion level of translator cascades

For a single bimolecular reversible toehold exchange reaction $X + F \rightleftharpoons Y + W$, where $X$ is the input signal, $F$ is the fuel, $Y$ is the displaced strand of this reaction and $W$ is the waste species. Assuming that the two toeholds in a toehold exchange reaction (i.e., toehold and balance domains in our nomenclature) have the same thermodynamic binding strength, the net reaction of a translator has $\Delta G^o \approx 0$ and the equilibrium constant of each reaction can be treated as 1. Thus for a single reaction, if the initial concentration for the reactants are $[X]_0 = \alpha$, $[F]_0 = 1$, at chemical equilibrium, the concentration of output strand $Y$ is $\frac{\alpha}{\alpha+1}$. This is obtained by solving the equilibrium concentrations: Assume the equilibrium concentration of $Y$ is $[Y]$, and the equilibrium concentration for the other species are $[W] = [Y], [X] = \alpha - [Y], [F] = 1 - [Y]$. According to the equilibrium expression for this reaction $K = \frac{[Y][W]}{[X][F]}$; in this case, $K = 1$. Solve the equation for $[Y]$, and we get $[Y] = \frac{\alpha}{\alpha+1}$.

Now we show the proof for the output concentration of a bimolecular (enthalpy-neutral) reversible reaction cascade.

**Theorem S4.1.** *(Theorem 1 in the main text) Given a linear enthalpy-neutral strand displacement cascades with $n$ ($n \geqslant 2$) layers, and with the fuel concentration 1 and the input concentration $\alpha$, the equilibrium concentration of the output signal is at least $\alpha\frac{1-2\alpha}{1-\alpha}$ independent of $n$.*

*Proof.* By conservation of mass, $[F_i] + [W_i] = 1$. Since $[W_i] \leqslant \alpha$ (we can't produce more waste than there was input), we get $[F_i] = 1 - [W_i] \geqslant 1 - \alpha$. Since the equilibrium constant is 1, at chemical equilibrium, for each reaction we have:

$$\frac{[X_{i+1}]}{[X_i]} = \frac{[F_i]}{[W_i]} \geqslant \frac{1-\alpha}{\alpha}.$$

Letting $\beta = \frac{1-\alpha}{\alpha} = \frac{1}{\alpha} - 1$, we get $[X_{i+1}] \geqslant \beta[X_i]$. This is a lower bound for $[X_{i+1}]$. Thus we have $[X_i] \leqslant \beta^{-(n+1-i)}[X_{n+1}]$.

Since the total concentration of all signal strands is conserved, we have:

$$[X_1] + [X_2] + \cdots + [X_i] + \cdots + [X_{n+1}] = \alpha$$

$$(\beta^{-n} + \beta^{-(n-1)} + \cdots + \beta^{-(n+1-i)} + \cdots + 1) \cdot [X_{n+1}] \geqslant \alpha$$

Since $\sum_{i=-n}^{0} \beta^i = \beta^{-n}\frac{1-\beta^{n+1}}{1-\beta}$, the above equation can be simplified as

$$[X_{n+1}] \cdot \beta^{-n}\frac{1-\beta^{n+1}}{1-\beta} \geqslant \alpha.$$

Thus the concentration of $X_{n+1}$ is

$$[X_{n+1}] \geqslant \frac{\alpha\beta^n(1-\beta)}{1-\beta^{n+1}} = \frac{\alpha(\frac{1}{\alpha}-1)^n(2-\frac{1}{\alpha})}{1-(\frac{1}{\alpha}-1)^{n+1}} = \frac{\alpha(2-\frac{1}{\alpha})}{\frac{1}{(\frac{1}{\alpha}-1)^n} - (\frac{1}{\alpha}-1)} = \frac{\alpha(\frac{1}{\alpha}-2)}{\frac{1}{\alpha}-1-\frac{1}{(\frac{1}{\alpha}-1)^n}} \geqslant \frac{\alpha(\frac{1}{\alpha}-2)}{\frac{1}{\alpha}-1} = \alpha\frac{1-2\alpha}{1-\alpha}.$$

□

### S5 Proofs for kinetic properties

#### S5.1 Toehold occlusion

**Theorem S5.1.** *(Theorem 2 in the main text) Toehold occlusion is not possible in a linear enthapy-neutral strand displacement cascades if and only if $N$ is a multiple of* shift*.*

*Proof.* Toeholds are occluded when overhangs can bind to them. The toehold of fuel $i$ lies at position $i \cdot \mathit{shift}$. Overhangs of fuel $i$ lie at positions between $i \cdot \mathit{shift} + N + 1$ and $i \cdot \mathit{shift} + N + \mathit{shift} - 1$. The claim follows by Lemma S2.2 ($p = \mathit{shift}$, $q = 0$ and $r = N + 1$). □

#### S5.2 Spurious strand displacement

##### S5.2.1 Spurious displacement in the absence of input

We begin by looking at spurious displacement of bottom domains which can only occur in toehold positions as these are the only positions with bottom domains in excess. We define *left flank* as the domain that is one unit to the right of a toehold domain.

**Observation S5.2.** *In the $i^{th}$ fuel, the left flank domain lies at the position $i \cdot \text{shift} + 1$.*

**Lemma S5.3.** *In the absence of input, spurious displacement of bottom domains in a linear enthalpy-neutral strand displacement cascade of arbitrary depth is possible (i) in left flank domains if and only if* $\text{shift} = 1$ *(i.e. the toehold domain can invade the left flank domain, see Fig. 2), and (ii) in balance domains if and only if $N$ is a multiple of* shift*.*

*Proof.* (i) By Observation S5.2 and Lemma S2.1, for a toehold position contains a left flank domain, set $p = \mathit{shift}$, $q = 0$ and $r = 1$ for their positions: $\forall i \in \mathbb{N}, \exists j \in \mathbb{N}$ such that $j \cdot p + q = i \cdot p + r$. This is possible if and only if $r - q$ is a multiple of $p$. It follows that $r - q = 1$ is a multiple of $p$, and thus $p = \mathit{shift} = 1$. (ii) By Observation S3.2 and Lemma S2.1, for a toehold position contains a balance domain, set $p = \mathit{shift}$, $q = 0$ and $r = N$ for their positions: $\forall i \in \mathbb{N}, \exists j \in \mathbb{N}$ such that $j \cdot p + q = i \cdot p + r$. It follows that $r - q = N$ is a multiple of $p = \mathit{shift}$. □

Now consider spurious displacement of top domains which can only occur in overhang positions as these are the only possible positions with top domains in excess.

**Lemma S5.4.** *In the absence of input, spurious displacement of top domains in a linear enthalpy-neutral strand displacement cascade of arbitrary depth is possible if and only if $N - 1$ is not a multiple of* shift*.*

*Proof.* By construction, when $\mathit{shift} = 1$ there are no overhang domains and therefore spurious displacement of top domains is not possible. Assume $\mathit{shift} > 1$.

By Lemma S3.5, it is not possible to spuriously displace top domains at balance positions. Thus any displacement of top domains of a fuel must be a proper prefix of its helix and therefore must include a left flank domain. By Observation S3.2, Observation S5.2 and Lemma S2.2, for overhang positions contains a left flank domain, set $p = \mathit{shift}$, $q = 1$ and $r = N + 1$ for their positions: $\forall i \in \mathbb{N}, \exists j \in \mathbb{N}$ such that $j \cdot p + q$ is contained in the interval $[i \cdot p + r,\ (i + 1) \cdot p + r - 2]$. This is possible if and only if $r - q - 1 > 0$ and $r - q - 1$ is not a multiple of $p$. It follows that $r - q - 1 = N - 1$ is not a multiple of $p = \mathit{shift}$. □

##### S5.2.2 Spurious displacement in the presence of input

A second type of spurious displacement is when a free signal strand (including the input), can act as a spurious invader of a fuel other than its designed target. In this case, particularly when the input concentration is significantly lower than fuel (as is typical), signal strands can become involved in numerous unproductive reactions, thus slowing (possibly significantly) signal propagation through every layer of the cascade.

**Lemma S5.5.** *Spurious displacement between signal strands and fuels is not possible in a linear enthalpy-neutral strand displacement cascade of arbitrary depth if and only if* shift $\geqslant N - 1$.

*Proof.* Domains of signal strand $i$ lie at positions between $i \cdot \textit{shift}$ and $i \cdot \textit{shift} + N - 1$. Signal strand $i$ is a spurious invader if it can displace any domains on some fuel $j > i$. Suppose signal strand $i$ is a spurious invader of fuel $j$; it must invade a prefix of fuel $j$'s double-stranded domains (its helix) which necessarily includes its left flank domain at position $j \cdot \textit{shift} + 1$ (Observation S5.2). It follows that $j \cdot \textit{shift} + 1 \leqslant i \cdot \textit{shift} + N - 1$, and thus $\textit{shift} \leqslant \frac{N-2}{j-i} \leqslant N - 2$ since $j > i$. Finally, suppose signal strand $i$ is not a spurious invader of any fuel $j > i$; then it cannot displace the left flank domain of fuel $j$, so $j \cdot \textit{shift} + 1 > i \cdot \textit{shift} + N - 1$ which implies $\textit{shift} > \frac{N-2}{j-i} = N - 2$ when $j = i + 1$. □

#### S6 Proofs for thermodynamic properties

All combinatorial arguments in this and the the following section are valid both in the single-molecule regime or when the fuels have multiple copies. Different fuels may also be present in different amounts.

The proofs in the following subsections are based on combinatorial arguments counting positions that have top or bottom domains in excess. Formally, we say a position is *top-limiting* if the number of bottom domains in that position is greater or equal to the number of top domains in that position. We say a position is *bottom-limiting* if the number of top domains in that position is greater or equal to the number of bottom domains in that position. The following observation follows directly from Lemma S3.4:

**Observation S6.1.** *In the absence of input: All toehold positions are top-limiting. All balance positions are top-limiting. All overhang positions are bottom-limiting.*

##### S6.1 Initial configuration is thermodynamically preferred without toehold occlusion

We first show that among the configurations with maximum bonds, the configurations with maximum separate complexes only contain complexes with exactly one top and one bottom strands. Then we show the proof for Theorem 4.

**Lemma S6.2.** *In the absence of input, in a maximum-bond configuration, every separate complex has at least one bottom strand and, for designs with* shift $\neq 1$*, every separate complex has at least one top strand.*

*Proof.* Since the configuration has the maximum number of bonds, the top domains in top-limiting positions must be bound and the bottom domains in bottom-limiting positions must be bound. Since every balance position is top-limiting and every top strand contains a balance domain, every top strand must bind with at least one bottom strand. For designs with *shift* $\neq 1$, every bottom strand contains a domain in an overhang position. Since every overhang position is bottom-limiting, every bottom strand must bind with at least a top strand. □

**Lemma S6.3.** *In the absence of input, in a maximum-bond configuration, for designs with* shift $= 1$*, any complex with two or more top strands must have two or more bottom strands.*

*Proof.* For designs with *shift* $= 1$, assume there exists a complex with two or more top strands and only one bottom strand. Let $N_b = N + 1$ be the number of bottom domains in this complex. Let $N_t \geqslant 2N$ be the number of top domains in this complex. For the configuration to have maximum bonds, every top domain needs to be bound. Thus $N_t \leqslant N_b$, and this implies that $N \leqslant 1$. However, recall from the definition of the design parameters in the main text that we require $N \geqslant 2$, contradicting that this design is valid. □

**Lemma S6.4.** *For designs without toehold occlusion, in the absence of input, any configuration that maximizes the number of separate complexes among the maximum-bond configurations consists of all complexes with exactly one top and one bottom strand.*

*Proof.* For designs with *shift* $\neq 1$, by Lemma S6.2, in a maximum-bond configuration every separate complex has at least one bottom strand and one top strand. Without input there is an equal number of top and bottom strands. Thus, the number of separate complexes is maximized when every complex has exactly one top strand and one bottom strand, as is the case in the initial configuration.

For designs with *shift* $= 1$, by Lemma S6.3 in a maximum-bond configuration every complex has a bottom strand. The initial configuration has maximum bonds, and it has as many complexes as bottom strands. Thus we know that any configuration that maximizes the number of separate complexes among the maximum-bond configurations must have complexes containing exactly one bottom strand (and $0$ or more top strands). If there exists a complex without top strands, then there must be another complex with at least two top strands, which implies it has at least two bottom strands (By Lemma S6.3). Thus we do not have two top stands in a complex and the only possibility is that every complex contains one top and one bottom strand. □

**Theorem S6.5.** *(Theorem 4 in the main text) For designs without toehold occlusion, given a linear enthalpy-neutral strand displacement cascade of arbitrary depth, in the absence of input, we have: (i) Among the configurations with maximum bonding, the initial configuration is the unique configuration that maximizes the number of separate complexes. (ii) The initial configuration has at least $N$ more bonds than any configuration with more separate complexes.*

*Proof.* (i) In the initial configuration, the $i$th top strand binds with the $i$th bottom strand. By Lemma S6.4, initial configurations have the maximum number of bonds. Since every complex has $N$ bonds, the total number of bonds in the maximum-bond configuration is $b \cdot N$, where $b$ is the number of bottom strands. Now we use contradiction to show that there does not exist any other configurations with the same number of bonds and the same number of separate complexes as the initial configurations. Suppose this configuration exists. By Lemma S6.4, every top strand only binds with one bottom strand. There must exist some bottom strand binding with one of its upstream top strands and some bottom strand binding with one of its downstream top strands. The number of bonds formed when a bottom strand binds with one of its upstream top strands is no more than $N$. The number of bonds formed when a bottom strand binds with one of its downstream top strands is smaller than $N$. Thus the total number of bonds formed in this configuration is smaller than $b \cdot N$. This contradicts with the assumption of maximum bonds.

As for (ii), the number of separate complexes in the initial configuration is equal to the number of bottom strands. For a configuration with more separate complexes, there must exist a complex with only top strands and no bottom strand. We consider two cases, *shift* $= 1$ and *shift* $\neq 1$. For *shift* $= 1$, since every position is top-limiting, every domain in the top strands must be bound. Considering that top strands do not bind each other, the complex with only top strands must have no bonds. Thus the configuration has at least $N$ fewer bonds than maximum bonds. For *shift* $\neq 1$, it directly follows from Theorem 6. Having a configuration with only a top strand is equivalent to having active output. A configuration with active output and $r$ bonds away from maximum has at least $\frac{N-r}{\mathit{shift}-1} - 1$ fewer separate complexes. For a configuration to have more separate complexes, $r$ needs to be at least $N$. □

#### S6.2 Enthalpic penalty to leak

To have an active output, it is necessary to break some bonds if there is no free top domain among the output region. The top-limiting positions do not have excess top domains, and thus these bonds must break. Therefore, it is sufficient to count the number of top-limiting positions among the output region. The following Lemma counts the number of balance and top-limiting positions in any output region. The Lemma is used in this subsection as well as in Lemma S6.10 in Section S6.3.

**Lemma S6.6.** *(i) Consider any output region. Its leftmost position is a toehold position, and among the rest there are* $\lfloor \frac{N-1}{\text{shift}} \rfloor$ *balance positions. (ii) In the absence of input, the number of top-limiting positions in any output region is* $\lceil \frac{N}{\text{shift}} \rceil$.

*Proof.* The leftmost position of an output region is a toehold position (by Observation S3.2). Here we count the balance positions among the rest of the output region (other than the leftmost). From Observation S3.2, balance positions are at $j \cdot \textit{shift} + N$ for $j = 0, \ldots, i$. We want to know how many of these lie in between $i \cdot \textit{shift} + 1$ to $i \cdot \textit{shift} + N - 1$: $i \cdot \textit{shift} + N - 1 \geqslant j \cdot \textit{shift} + N \geqslant i \cdot \textit{shift} + 1$. Thus there are $\lfloor \frac{N-1}{\textit{shift}} \rfloor$ balance positions among the rest of the output region.

To count the number of top-limiting positions among the output region, first we know that the leftmost position is top-limiting (by Observation S6.1). For the rest of the output region, some are overhang positions and some are balance positions (by Lemma S3.5). By Observation S6.1, the total number of top-limiting positions among the active output is $\lfloor \frac{N-1}{\textit{shift}} \rfloor + 1 = \lceil \frac{N}{\textit{shift}} \rceil$. □

**Theorem S6.7.** *(Theorem 5 in the main text) Given a linear enthalpy-neutral strand displacement cascade of arbitrary depth, in the absence of input, any configuration having active output has at least* $\lceil \frac{N}{\text{shift}} \rceil$ *fewer bonds than the maximum-bond configuration.*

*Proof.* For a configuration to have active output, the bonds in the top-limiting positions among the output region must break, thus it is sufficient to count the number of top-limiting positions. By Lemma S6.6, the configuration with active output has at least $\lceil \frac{N}{\textit{shift}} \rceil$ fewer bonds than the maximum-bond configuration. □

##### S6.3 Entropic penalty to leak

For designs without toehold occlusion, we have the following observations for the position types:

**Observation S6.8.** *For designs without toehold occlusion, toehold positions and overhang positions do not overlap, while all toehold and balance positions overlap.*

To count the difference in the number of separate complexes between the thermodynamically preferred configuration and the active-output configuration, it is useful to count the number of bottom domains in overhang positions first.

**Lemma S6.9.** *For designs without toehold occlusion, a bottom strand contains* $N - \frac{N}{\text{shift}}$ *overhang positions, and a top strand contains* $\text{shift} - 1$ *more overhang positions compared to a bottom strand.*

*Proof.* A bottom strand has $N + 1$ domains and the distance between two toehold position is *shift*. Thus a bottom strand contains $\lfloor \frac{N+1}{shift} \rfloor + 1$ toehold positions. By Observation S6.8, the number of overhang position in a bottom strand is $N + 1 - (\lfloor \frac{N+1}{shift} \rfloor + 1) = N - \frac{N}{shift}$ ($N$ is a multiple of *shift*, by Theorem 2). Since there is no toehold occlusion, an overhang position does not contain any toeholds. A top strand has $shift - 1$ (by Observation S3.1) more overhang positions than a bottom strand. □

**Lemma S6.10.** *For designs with* $shift \neq 1$ *and no toehold occlusion, in the absence of input, if a configuration with active output and* $r$ ($r \in \mathbb{N}$, $r \geqslant \frac{N}{\text{shift}}$) *bonds away from maximum, there are* $r - \frac{N}{\text{shift}}$ *unbound bottom domains in overhang positions.*

*Proof.* By Lemma S6.6 and Observation S6.8, the configuration with active output contains $\frac{N}{shift}$ balance positions among the open positions in active output. Since balance positions are top-limiting (Observation S6.1), the number of unbound bottom domains in balance positions among the open positions in active output is equal to $\frac{N}{shift}$. Thus the remaining budget of unbound bottom domains for overhang positions is $r - \frac{N}{shift}$. □

For the purpose of the arguments in this section, we define *normal form* as: Given a linear enthalpy-neutral strand displacement cascade of arbitrary depth, and no input signal, a complex is normal form if the number of top strands is equal to the number of bottom strands. A configuration is normal form if every complex is normal form.

**Lemma S6.11.** *For designs with* $shift \neq 1$ *and no toehold occlusion, in the absence of input, if a configuration with* $r$ ($r \in \mathbb{N}$, $r \geqslant \frac{N}{\text{shift}}$) *bonds away from maximum is not normal form, then some complex has at least* $\frac{N-r}{\text{shift}-1}$ *top strands.*

*Proof.* Since the configuration is not normal form, there must exist a complex $P$ with more bottom strands ($b$) than top strands ($t$). Since $b > t$, $b \geqslant t + 1$. Let $N_t$ be the number of overhang positions in a top strand, and $N_b$ be the number of overhang positions in a bottom strand ($N_b = N - \frac{N}{shift}$, Lemma S6.9). By Observation S6.1, all overhang positions are bottom-limiting, and thus in maximum-bond configuration, all bottom domains in overhang positions must be bound. In a configuration with $r$ bonds away from maximum bonds, the number of bound bottom domains in overhang positions is $N_b \cdot b - (r - \frac{N}{shift})$ (Lemma S6.10). The number of top domains in overhang positions is $N_t \cdot t$. Thus we know that $N_t \cdot t \geqslant N_b \cdot b - (r - \frac{N}{shift}) \geqslant N_b \cdot (t + 1) - (r - \frac{N}{shift})$. By Lemma S6.9, $N_t - N_b = shift - 1$. Simplifying the inequality we get $t \cdot (shift - 1) \geqslant N_b - (r - \frac{N}{shift})$, and thus $t \geqslant \frac{N-r}{shift-1}$. □

**Lemma S6.12.** *For designs with shift $\neq 1$ and no toehold occlusion, in the absence of input, if a configuration with $r$ ($r \in \mathbb{N}$, $r \geqslant \frac{N}{\text{shift}}$) bonds away from maximum is normal form and a complex has an active output, then some complex has at least $\frac{N-r}{\text{shift}-1}$ top strands.*

*Proof.* Since the configuration is normal form, the active output complex $P$ has equal top ($t$) and bottom ($b$) strands ($t = b$). Let $N_t$ be the number of overhang positions in a top strand, and $N_b$ be the number of overhang positions in a bottom strand ($N_b = N - \frac{N}{shift}$, Lemma S6.9). Among the positions for the $N$ free domains in active output complex $P$, the number of overhang positions is the same as the number of overhang positions in a bottom strand $N_b$. By Observation S6.1, all overhang positions are bottom-limiting, and thus in the positions for the $N$ free domains in $P$, there must be $N_b$ free top domains from overhang positions. Since the configuration is $r$ bonds away from maximum bonds, some free top domains can be produced by breaking bonds. The number of top domains in the overhang positions for the $N$ free domains in active output complex $P$ should be at least $N_b - (r - \frac{N}{shift})$ more than bottom domains (Lemma S6.10). We know that the difference between the total number of top and bottom domains in overhang positions in $P$ is $t \cdot (shift - 1)$ (Lemma S6.9). Thus $t \cdot (shift - 1) \geqslant N_b - (r - \frac{N}{shift})$. Simplifying the inequality we get $t \geqslant \frac{N-r}{shift-1}$. □

**Theorem S6.13.** *(Theorem 6 in the main text) For designs with* shift $\neq 1$ *and no toehold occlusion, given a linear enthalpy-neutral strand displacement cascade of arbitrary depth, in the absence of input, any configuration with active output and $r$ ($r \in \mathbb{N}, r \geqslant \frac{N}{\text{shift}}$) bonds away from maximum has at least $\frac{N-r}{\text{shift}-1} - 1$ fewer separate complexes than the initial configuration.*

*Proof.* By Lemma S6.11 and S6.12, for a configuration with active output, there must exist some complex with at least $\frac{N-r}{shift-1}$ top strands. Thus the entropic penalty to leak is $\frac{N-r}{shift-1} - 1$. □

### S7 Reconfiguration

In this section, we prove that in certain cases the hard-to-reverse displacement descriebed in the main text is thermodynamically penalized by connecting it to active output. Unfortunately, our argument holds only when *shift* is large leading to a small thermodynamic penalty; obtaining the penalty for arbitrary *shift* remains an area for further research.

For the purpose of the arguments, we use index ($i \in \mathbb{N}$) to refer the top or bottom strands in the $i$th fuel in the initial configuration. As before, we allow multiple copies of all the strands, as long as there are exactly as many top strands with index $i$ as bottom strands with index $i$. Given a linear enthalpy-neutral strand displacement cascade of arbitrary depth, and no input signal, we say a configuration is *unmatched* if there is an index $i$ and a complex $P$ such that $P$ contains a different number of top and bottom strands with index $i$. We say $P$ is *top-heavy* at the index $i$ if there are more top strands with index $i$ in $P$. Similarly, $P$ is *bottom-heavy* if there are more bottom strands with index $i$ in $P$.

Unmatched configurations exactly capture hard-to-reverse reconfiguration discussed in the main text. If the configuration is *matched*, then there is a sequence of (fast) unimolecular reactions that leads to the initial configuration: we can split any complex into initial fuel complexes. On the other hand, no sequence of unimolecular reactions leads to the initial configuration from an unmatched configuration and (slow) bimolecular reactions are necessary to undo the spurious displacement.

Our goal is to argue that unmatched configurations are thermodynamically penalized, and thus any sequence of spurious displacement reactions leading to them is unfavorable. First, we note that any design without toehold occlusion has an unmatched configuration with maximum bonds (see Figure S1 for an example). Thus, we do not have a purely enthalpic penalty as we do for active output (Theorem 5).

**Observation S7.1.** *For designs without toehold occlusion, given a linear enthalpy-neutral strand displacement cascade of arbitrary depth, in the absence of input, there exists an unmatched maximum-bond configuration.*

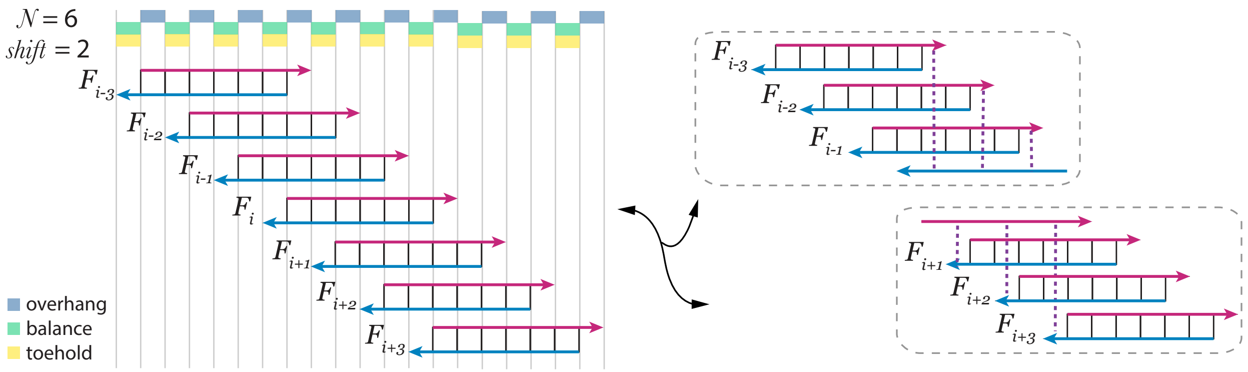

**Figure S1:** An example of an unmatched configuration for design $N = 6$, *shift* $= 2$ with maximum bonds. A similar configuration, in which the top and bottom strands of an initial fuel complex are separated without net loss of bonds, exists for all designs without toehold occlusion.

While unmatched configurations may not incur any enthalpy penalty, they may be penalized entropically. For example, in Figure S1, the unmatched configuration on the right has five fewer separate complexes than the initial configuration on the left. Indeed, we now show that for certain choices of *shift* unmatched configurations provably incur an entropic thermodynamic penalty compared with the initial configuration. We do this by making a connection between unmatched configurations and configurations with active output, which we already know incur an entropic penalty.

**Lemma S7.2.** *Assume* shift $> \frac{N}{2}$*. Suppose a complex $P$ is top-heavy at index $i$ and $P$ does not contain any bottom strands with index larger than $i$. Then $P$ has active output.*

*Proof.* We know that the balance domain of the bottom strand at index $(i-1)$ lies at position $(i-1) \cdot \textit{shift} + N$. The active output of the $i$th top strand starts at position $(i+1) \cdot \textit{shift}$. Since $\textit{shift} > \frac{N}{2}$, $(i-1) \cdot \textit{shift} + N = i \cdot \textit{shift} + N - \textit{shift} < i \cdot \textit{shift} + 2\textit{shift} - \textit{shift} = i \cdot \textit{shift} + \textit{shift}$, there must exist an active output in the $i$th top strand regardless of whether there are more bottom strands at index $k$ ($k < i$) in $P$. □

**Lemma S7.3.** *For designs with* shift $> \frac{N}{2}$*, given a linear enthalpy-neutral strand displacement cascade of arbitrary depth, in the absence of input, for any unmatched configuration $C$, there is an active output configuration $C'$ and $d$ ($d \in \mathbb{N}$), such that $C'$ has $d$ more separate complexes but at most $d$ fewer bonds than $C$.*

*Proof.* Let $i$ be the *largest* index for which there is a top-heavy complex at index $i$; call this complex $P$. Note that $P$ can't be bottom-heavy for any index $j > i$ since that would imply that there is another complex that is top-heavy at index $j > i$, contradicting our choice of $i$. Thus for all $j > i$, complex $P$ contains the same number of top and bottom strands of index $j$ (which could be zero count).

If $P$ does not contain any strands of index larger than $i$ then this lemma follows with $d = 0$ by Lemma S7.2. Otherwise, let $j > i$ be the largest index that $P$ contains. since $j$ is the largest index in $P$, the overhangs of the top strands with index $j$ are not bound in $P$. Thus we can split off pairs of top and bottom strands of index $j$, possibly breaking the bond of the toehold of the bottom $j$ strand with some other top strand in $P$. Each such split increases the number of separate complexes by 1 and decreases the number of bonds by at most 1. We can repeat the process until $i$ is the largest index that $P$ contains. Then this lemma follows by Lemma S7.2, with $d$ being the total number of pairs we split off. □

Unfortunately, because of the restriction $\textit{shift} > \frac{N}{2}$, using Theorem 6 yields only weak results. Note that the only such design without toehold occlusion is $\textit{shift} = N$. Further, the entropic penalty is small for large *shift*. Thus, combining Lemma S7.3 and Theorem 6 only shows that there is an entropy penalty of 1 for $N = \textit{shift}$ when $r = 0$.

#### S8 Comparing leak reduction in enthalpy-neutral and NLD designs

Recently, a leak reduction method ("*NLD* design") was proposed where leak requires binding of $N$ separate fuels, where the choice of $N$ can be made arbitrarily large [2, 3]. The chance of this happening decreases exponentially with $N$; equivalently, the energy barrier to leak can be made arbitrarily large by means of the entropy component.

Compared with the *NLD* design, we achieve a stronger thermodynamic guarantee of correct behavior in the *EN* design. In the *NLD* design, a leaked upstream signal can start a cascade which gains enthalpy from toehold binding for every downstream strand displacement step, which could be overall favorable for long cascades. Thus, the prior work corresponds to a kinetic barrier in our model. In contrast, in our *EN* design, a leaked upstream signal cannot be driven by the enthalpy of forming new bonds downstream. In fact, leak requires the overall *loss* of bonds in our design (enthalpic penalty), and leak in our designs is always thermodynamically penalized.

This is not, by itself, enough to necessarily prefer the *EN* designs to *NLD* designs because a high kinetic barrier might practically prevent leak better than a relatively low thermodynamic penalty. Thus, we want to quantitatively compare the two. Both the *NLD* kinetic barrier and the enthalpy-neutral thermodynamic penalty can be made arbitrarily large by proper choice of parameters, so we compare the size of both as a common function of the complexity of the system as captured by the number of fuel complexes and the total amount of DNA.

In the following subsections, we show that for the same number of fuel complexes, our end-state thermodynamic penalty to leak is $\beta + 1$ the size of the *NLD* kinetic barrier when the free energy of forming a new bond (toehold) is $\beta$ times as favorable as a new separate complex. Thus our penalty to leak is always larger, and that when enthalpy and entropy are balanced [1] ($\beta = 1$) our penalty is twice as high. For larger $\beta$ (e.g., especially high concentrations), the advantage of our design is even larger (any *shift*). Designs with small *shift* are also more efficient in terms of total DNA used.

Note that if the clamps in the *NLD* design are extended to toehold size, the *NLD* design can be categorized into one class of the *EN* design. For example, the *SLD* design would be categorized as $N = 4$ and *shift* $= 3$, and the *DLD* design would be categorized as $N = 6$ and *shift* $= 3$. According to the taxonomy developed here, this extended *NLD* design for any redundancy $N_{NLD}$ has toehold occlusion and spurious strand displacement. In this sense, the *EN* design is broader than the *NLD* design with toehold-size clamps, allowing for more flexibility in balancing desired and undesired properties.

##### S8.1 Recap of the energy penalty to leak in the *NLD* and *EN* designs

For a fair comparison, we focus on a full translator which generates an output of independent sequence from the input. Let $N_{NLD}$ be the number of "long domains" (as defined in [3, 2]); this is also the number of fuels in a translator in the *NLD* design. Note that $N_{NLD}$ does not directly corresponds to our parameter $N$ as our domains are toehold-size.

The *NLD* kinetic barrier to active output (leak) is due to reducing the number of separate complexes (entropic barrier). Specifically, the entropic penalty to leak for the *NLD* design is $N_{NLD} - 1$ entropy units [2]. (This acts as a kinetic barrier: although additional separate complexes cannot be

[1] It is experimentally desirable to be close to the regime where forming a new bond (toehold) or a new separate complex is equally favorable. This regime balances high concentration (reaction speed) with toehold occlusion and spurious signal-fuel binding.

generated, the downstream displacement of a leaked signal gains enthalpy from toehold binding for every downstream strand displacement step.)

In contrast, for our enthalpy-neutral strand displacement design there is a combined entropic and enthalpic penalty to leak, that is not transient. Specifically, with *shift* $\neq 1$ and no toehold occlusion ($N$ is a multiple of *shift*, Theorem 2), the enthalpic penalty to leak is at least $\frac{N}{shift}$ (Theorem 5). Rephrasing Theorem 6, the entropic penalty to leak is at least

$$\frac{N - \frac{N}{shift} - r'}{shift - 1} - 1$$

where $r'$ is the number of *additional* bonds lost over the minimum of $\frac{N}{shift}$.

##### S8.2 Enthalpy equals entropy regime

Consider first the regime where forming a new bond (toehold) or a new separate complex is equally favorable— this regime balances high concentration (reaction speed) with toehold occlusion and spurious signal-fuel binding. (We consider other regimes below.) Since a unit of enthalpy and a unit of entropy are then equal, we can simply add up enthalpic and entropic contributions to the leak penalty, defining a common free-energy *unit*. The height of the kinetic barrier to leak in the *NLD* design has the units:

$$E_{NLD} = N_{NLD} - 1$$

For our enthalpy-neutral design, summing up the enthalpic and entropic penalties to leak we get the total units:

$$E = (\frac{N}{shift} + r') + (\frac{N - \frac{N}{shift} - r'}{shift - 1} - 1) \geqslant 2\frac{N}{shift} - 1$$

Thus, we can say that the total energy penalty to leak of the enthalpy-neutral strand displacement design is $E = 2\frac{N}{shift} - 1$ units.

We can now quantitatively compare the *NLD* barrier and the enthalpy-neutral design penalty, keeping the number of fuels constant. The number of fuels for a translator in the *NLD* design is $N_{NLD}$, and for the enthalpy-neutral design it is $\frac{N}{shift}$ (Observation S3.3). Thus for the same number of fuels, the enthalpy-neutral design penalty to leak has asymptotically twice the height.

We now focus on another natural measure of system complexity, the total amount of DNA needed to achieve $E = E_{NLD}$. How can we compare toehold-size domains and long domains in terms of total DNA? A reasonable assumption is that a complex bound with two or more toehold-size domains will not dissociate; thus we can estimate that an *NLD* "long" domain corresponds to $l > 1$ domains in our design. This allows us to measure the amount of total DNA in both systems using a common unit of (toehold-size) *domains*.

First we count the number of domains in a translator in the *NLD* design. A translator has $N_{NLD}$ fuels. Each fuel has $N_{NLD}$ bound long domains, 1 unbound long domain as overhang and 1 unbound domain as toehold. Thus the total number of domains in a translator in the *NLD* design is

$$D_{NLD} = N_{NLD} \cdot (2N_{NLD} \cdot l + l + 1) = 2l \cdot N_{NLD}^2 + l \cdot N_{NLD} + N_{NLD}$$

Assuming $E_{NLD} = E$, we have $N_{NLD} = 2\frac{N}{shift}$. Therefore,

$$D_{NLD} = \frac{8l}{shift^2}N^2 + \frac{2l + 2}{shift}N$$

Now we count the number of domains in a translator in the enthalpy-neutral strand displacement design. A translator has $\frac{N}{\mathit{shift}}$ of fuels. Each fuel has $N$ bound domains, $\mathit{shift} - 1$ unbound domains as overhang and $1$ unbound domain as toehold. So the total number of domains in an enthalpy-neutral translator is

$$D = \frac{N}{\mathit{shift}} \cdot (2N + \mathit{shift} - 1 + 1) = \frac{2}{\mathit{shift}} N^2 + N$$

Depending on the choice of *shift* and $l$, the total amount of DNA between these two designs varies with a threshold at

$$l_0 = \frac{\mathit{shift}(2N + \mathit{shift} - 2)}{2(4N + \mathit{shift})}$$

When $l \geqslant l_0$, $D_{NLD} \geqslant D$; and when $l \leqslant l_0$, $D_{NLD} < D$. Thus, for small *shift* (less than 8 assuming $l = 2$), the enthalpy-neutral design achieves the same height energy penalty as the *NLD* energy barrier using less total DNA.

#### S8.3 Other regimes

Let $\beta$ be the relative weight of a unit of enthalpy (additional bond) compared with a unit of entropy (additional separate complex). For example, all else being equal, at high concentrations we can expect $\beta > 1$ and at low concentrations $\beta < 1$. Given that the *NLD* leak barrier is entirely entropic, while the enthalpy-neutral leak penalty has enthalpic and entropic components, we expect the advantage of our enthalpy-neutral design to be greatest when $\beta > 1$. Indeed, we now argue that for any $\beta$, the enthalpy-neutral leak penalty can be made a factor of $\approx \beta + 1$ larger than the barrier in the *NLD* design using the same number of fuels.

Recall the entropic barrier in the *NLD* design is: $E_{NLD} = N_{NLD} - 1$ *entropy units*. For our enthalpy-neutral design, converting the enthalpy part of the penalty to entropic units, we obtain a combined energy penalty in entropic units:

$$E = \beta(\frac{N}{\mathit{shift}} + r') + (\frac{N - \frac{N}{\mathit{shift}} - r'}{\mathit{shift} - 1} - 1) = \beta\frac{N}{\mathit{shift}} + \frac{N - \frac{N}{\mathit{shift}}}{\mathit{shift} - 1} - 1 + r'(\beta - \frac{1}{\mathit{shift} - 1})$$

We can use *shift* large enough that $\beta \geqslant \frac{1}{\mathit{shift}-1}$. Then the above expression is minimized when $r' = 0$, and so we can say that the enthalpy-neutral energy penalty to leak is $E = \beta\frac{N}{\mathit{shift}} + \frac{N - \frac{N}{\mathit{shift}}}{\mathit{shift}-1} - 1 = (\beta + 1)\frac{N}{\mathit{shift}} - 1$.

We are interested in the ratio $E/E_{NLD}$ when the number of fuels is the same for the two designs. Recall that the number of fuels for a translator in both designs is the same when $N_{NLD} = \frac{N}{\mathit{shift}}$. Since $E_{NLD} = N_{NLD} - 1$ and $E = (\beta + 1)\frac{N}{\mathit{shift}} - 1$, we have $E/E_{NLD} \approx \beta + 1$.

#### S9 Orthogonal toeholds

Our argument relies on the strong assumption that all domains are orthogonal. In reality, given the limited size of a domain (that of a toehold), as the number of distinct domains increases, it is not possible to make all the domains orthogonal. Nonetheless we note that in certain cases having the same sequence in multiple domain positions seems to pose no problem. (For example, in designs where $N$ is not a multiple of *shift*, all the balance domains could have the same sequences.) Future work could further explore how to assign the same domains without undesired interactions and how the number of orthogonal domains needed scales with the length of double-stranded region $N$.

#### S10 Sequence design

The sequences for the typical leaky translator ($SLD$) and the previously reported leak reduction method based exclusively on an entropic barrier ($DLD$) are from reference [3].

All other sequences were generated using the following method. The sequence space of the linear strand displacement cascade is represented by a row of unique domains. First, we generated a pool of unique domains. Each domain has 7 nucleotides with equal probability of $A$, $T$, $G$. The 3-letter $ATG$ alphabet was used for all bottom strands (and naturally the complementary $ATC$ alphabet for the top strands) in order to minimize self-complementarity of a strand (no possibility to form strong $C$-$G$ bonds) [4, 5]. Every domain contains $2$ or $3$ $G$s and no $4$ consecutive $A$'s or $T$'s. In addition, domains in the pool have Hamming Distance greater than $2$. Then we drew domains from the pool and assembled a single contiguous sequence as the sequence space candidate. The candidates were selected if they do not contain more than $4$ of the same nucleotide in a row, and do not contain $7$ $A/T$'s in a row. Among all the candidate sequences, we chose the sequence with the minimal length of the longest repeated segments. Mathematica code written to generate sequences according to these steps is available upon request.

Due to the enthalpic contribution from the fluorophore and quencher interaction, the balance domains in reporters were shortened to 5 bases rather than 7.

#### S11 Data fitting and normalization

A calibration curve was used to convert arbitrary fluorescence units to the corresponding signal concentration. We obtained different calibration curves for each reporter. The data in the following figures were normalized by calibration curves: Figure 6, positive triggering signal in Figure S4 (this was used to compute the signal-to-leak ratio shown in Figure 8b inset) and Figure 9.

The reporter was titrated with the top strand of its upstream fuel at different concentrations (i.e., input signal strand to the reporter). The average of the last ten data points from 4-hour kinetics were calculated as the signals at reaction completion (although reactions reach completion at about half an hour). The calibration curve was made by fitting the triggered signals to the initial concentrations of invading top strands through the model

$$top + reporter \overset{K}{\rightleftharpoons} reporter.top + waste$$

($K$ is the equilibrium constant), also shown in Figure S2a.

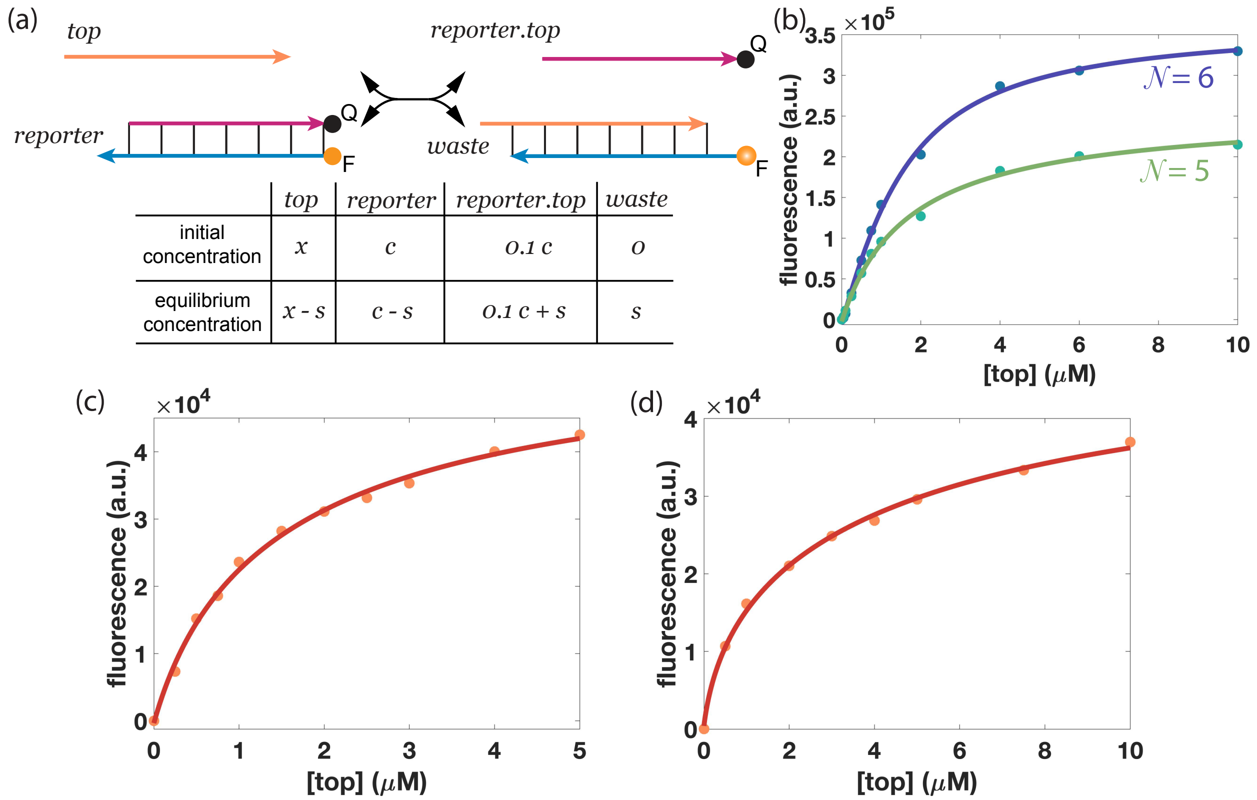

**Figure S2:** The model used for concentration calibration. (a) One-step reaction model of reporter triggering. The top strand of the reporter is labeled with a quencher (Q). The bottom strand of the reporter is labeled with a fluorophore (F). (b) The calibration curves for reporters with $N = 6$ and $N = 5$ (Figure 6). The nominal initial concentration of reporters was 2 $\mu$M. (c) The calibration curve for the reporter with $N = 5$, *shift* $= 3$ (Figure 9). The nominal initial concentration of reporter was 5 $\mu$M. (d) The calibration curve for the reporter of the enthalpy-neutral strand displacement design shown in Figure S4 (experiment comparing the enthalpy-neutral strand displacement design with the prior *NLD* design). The nominal initial concentration of reporter was 6 $\mu$M.

Assume the initial concentrations of the invading top strand (top) is $x$, and the initial concentration of the reporter is $c$. Since the reporter was prepared with $10\%$ excess of its top strand (reporter.top), the initial concentration of reporter.top is $0.1c$. At equilibrium, assume the waste concentration is $s$. Then the equilibrium concentration of other species can be derived accordingly and they are listed in Figure S2a. The concentrations satisfy

$$K = \frac{[\textit{waste}][\textit{reporter.top}]}{[\textit{top}][\textit{reporter}]} = \frac{s \cdot (0.1c + s)}{(x - s)(c - s)}$$

Simplifying the expression for $s$, we get:

$$s = \frac{0.1c + x \cdot K + c \cdot K + \sqrt{4x \cdot c \cdot K \cdot (1 - K) + (0.1c + x \cdot K + c \cdot K)^2}}{2(K - 1)} \tag{1}$$

The waste species is the only species emitting fluorescence signal and the fluorescence signal is linearly dependent on the waste species' concentration with the linear parameter $\Delta F$. Assume the fluorescence signal is $F$ when the waste species concentration is $s$. We have

$$F = F_0 + \Delta F \cdot s \tag{2}$$

where $F_0$ is the background signal reported by the instrument.

For each titration experiment, with a set of known $x$ and the collected signal $F$, we fit them to equations (1) and (2) and get a calibration curve with fitted parameters $c$, $K$, $\Delta F$ and $F_0$. We noticed that the fitted parameter $c$ sometimes differs from the nominal concentration used in experiments: Although the initial concentration of the reporter $c$ is known, fixing the parameter $c$ may not give a good fit especially when the initial concentration of the reporter is high (e.g. $5\ \mu M$). Thus model here is used as the phenomenological model instead of the mechanistic model. Figure S2 shows the calibration curves used in this paper.

In experiments generating a relatively small amount of output, we expect the fluorescence signal to be linear. Thus, the following data were normalized to a linear calibration curve: experiments comparing designs with different energy penalty to leak at different temperatures (Figure 5, Figure S3), and experiments comparing the leak at equilibrium between the enthalpy-neutral strand displacement design (leak in Figure 8b) and the *NLD* design from prior work.

### S12 Triggered signals for designs with different energy penalty to leak

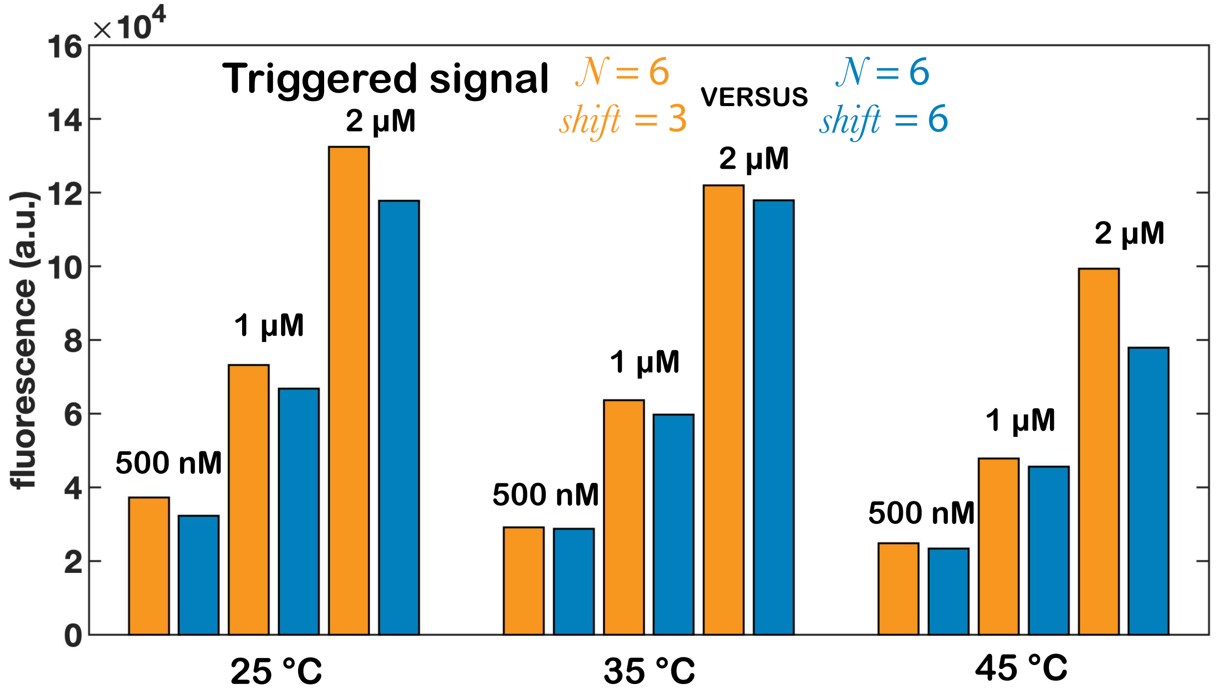

**Figure S3:** Comparison of desired triggering of designs from Figure 5. Although the $N = 6$, *shift* $= 3$ design had significantly less leak as shown in Figure 5, it had similar (or even slightly higher) completion level. These two designs share the same reporter. Fluorescence levels were measured at different thermodynamic equilibria (25 °C, 35 °C and 45 °C) after annealing with the input strand. The concentrations for fuels and reporters are labeled in the figure. The input strands were added at half the concentration of fuels.

#### S13 Comparison of triggered signals between the prior *NLD* designs and the enthalpy-neutral strand displacement design

To compare the signal-to-leak ratio between the two designs, we need to know the triggered signals and the leak concentrations. Leak concentrations are shown in Figure 8. As shown in Figure S4, the triggered signals for the enthalpy-neutral strand displacement design were measured and normalized to the calibration curve in Figure S2d.

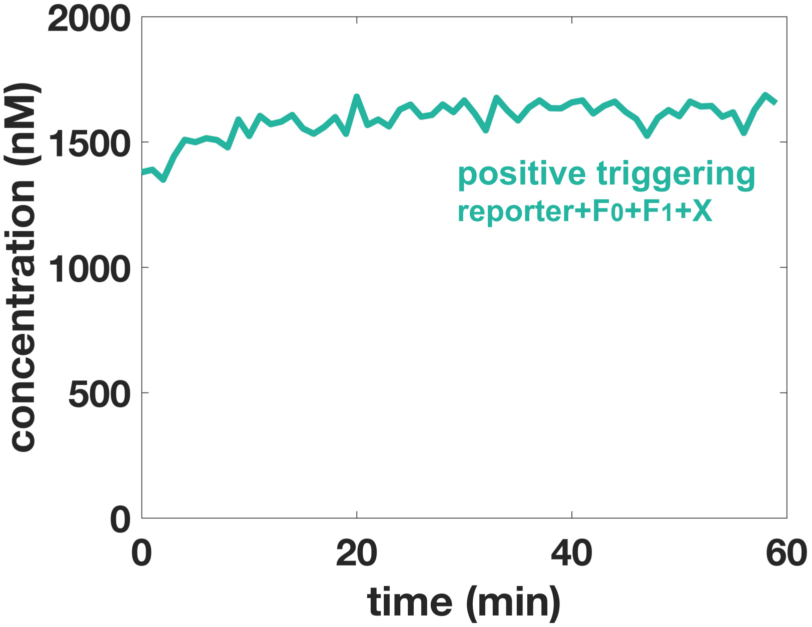

**Figure S4:** Kinetics of the triggered signal for the enthalpy-neutral strand displacement design. The concentrations for fuels and input were 5 $\mu$M, and the reporter was 6 $\mu$M. The reaction temperature was 25 °C.

In Figure 8, the triggered signals for the *NLD* designs were assumed to reach full completion. For example, in the case when the fuels and input were 5 $\mu$M, and the reporter was 6 $\mu$M, the triggered signal was assumed to reach 5 $\mu$M.

#### S14 Sequences

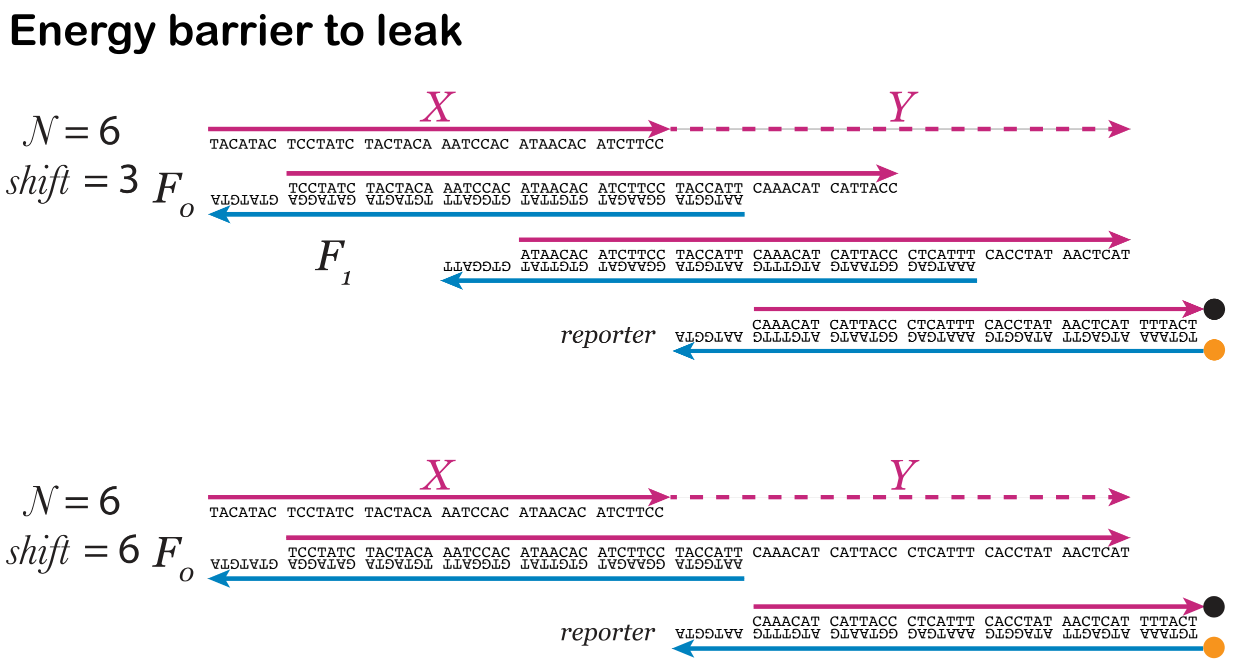

**Figure S5:** Sequences used for the experiments comparing the designs with different energy penalty to leak in Figure 5.

| | |
|---|---|
| N6.X | TACATAC TCCTATC TACTACA AATCCAC ATAACAC ATCTTCC |
| N6S3.F0.top | TCCTATC TACTACA AATCCAC ATAACAC ATCTTCC TACCATT CAAACAT CATTACC |
| N6S3.F0.bottom | AATGGTA GGAAGAT GTGTTAT GTGGATT TGTAGTA GATAGGA GTATGTA |
| N6S3.F1.top | ATAACAC ATCTTCC TACCATT CAAACAT CATTACC CTCATTT CACCTAT AACTCAT |
| N6S3.F1.bottom | AAATGAG GGTAATG ATGTTTG AATGGTA GGAAGAT GTGTTAT GTGGATT |
| N6S6.F0.top | TCCTATC TACTACA AATCCAC ATAACAC ATCTTCC TACCATT CAAACAT CATTACC CTCATTT CACCTAT AACTCAT |
| N6S6.F0.bottom | share the same sequence as N6S3.F0.bottom |
| N6.Rep.top | CAAACAT CATTACC CTCATTT CACCTAT AACTCAT TTTACT/3IAbRQSp/ |
| N6.Rep.bottom | /56-ROXN/TGTAAA ATGAGTT ATAGGTG AAATGAG GGTAATG ATGTTTG AATGGTA |

#### Toehold occlusion

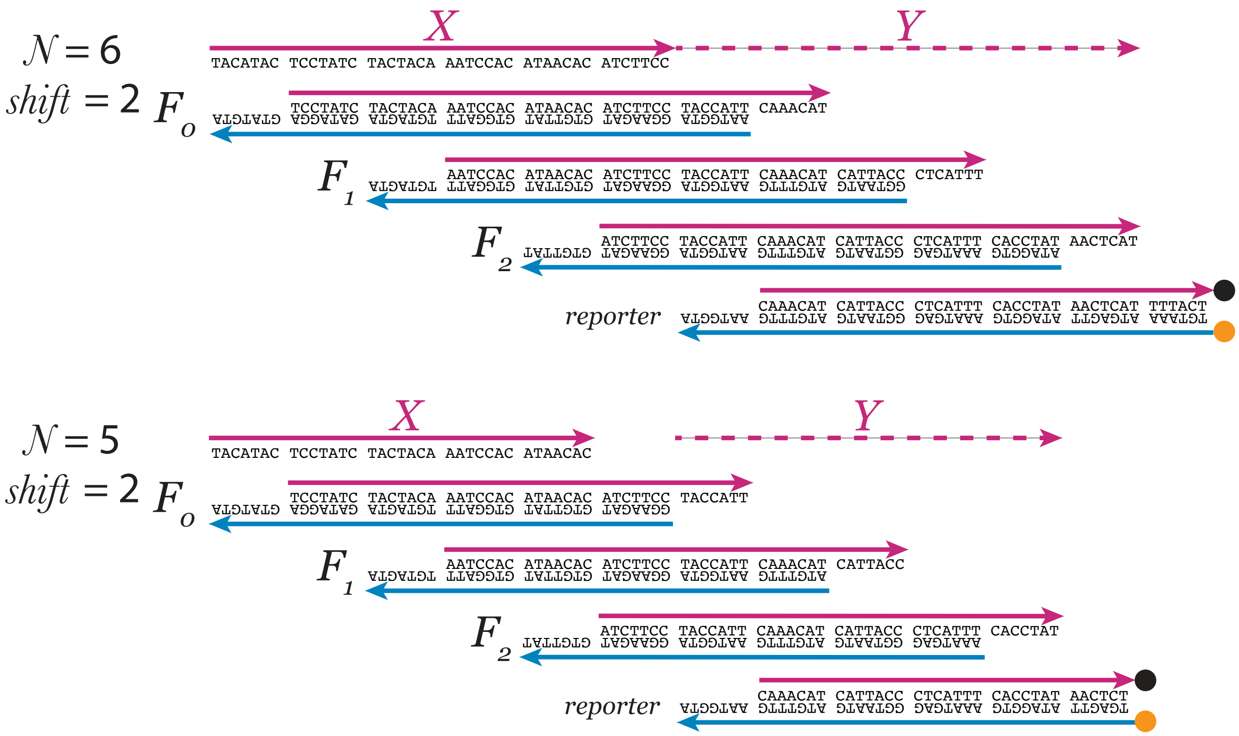

**Figure S6:** Sequences used for the experiments comparing the designs with and without toehold occlusion in Figure 6.

| | |
|---|---|
| N6.X | TACATAC TCCTATC TACTACA AATCCAC ATAACAC ATCTTCC |
| N6S2.F0.top | TCCTATC TACTACA AATCCAC ATAACAC ATCTTCC TACCATT CAAACAT |
| N6S2.F0.bottom | AATGGTA GGAAGAT GTGTTAT GTGGATT TGTAGTA GATAGGA GTATGTA |
| N6S2.F1.top | AATCCAC ATAACAC ATCTTCC TACCATT CAAACAT CATTACC CTCATTT |
| N6S2.F1.bottom | GGTAATG ATGTTTG AATGGTA GGAAGAT GTGTTAT GTGGATT TGTAGTA |
| N6S2.F2.top | ATCTTCC TACCATT CAAACAT CATTACC CTCATTT CACCTAT AACTCAT |
| N6S2.F2.bottom | ATAGGTG AAATGAG GGTAATG ATGTTTG AATGGTA GGAAGAT GTGTTAT |
| N5.X | TACATAC TCCTATC TACTACA AATCCAC ATAACAC |
| N5S2.F0.top | TCCTATC TACTACA AATCCAC ATAACAC ATCTTCC TACCATT |
| N5S2.F0.bottom | GGAAGAT GTGTTAT GTGGATT TGTAGTA GATAGGA GTATGTA |
| N5S2.F1.top | AATCCAC ATAACAC ATCTTCC TACCATT CAAACAT CATTACC |
| N5S2.F1.bottom | ATGTTTG AATGGTA GGAAGAT GTGTTAT GTGGATT TGTAGTA |
| N5S2.F2.top | ATCTTCC TACCATT CAAACAT CATTACC CTCATTT CACCTAT |
| N5S2.F2.bottom | AAATGAG GGTAATG ATGTTTG AATGGTA GGAAGAT GTGTTAT |
| N6.Rep.top | CAAACAT CATTACC CTCATTT CACCTAT AACTCAT TTTACT/3IAbRQSp/ |
| N6.Rep.bottom | /56-ROXN/TGTAAA ATGAGTT ATAGGTG AAATGAG GGTAATG ATGTTTG AATGGTA |
| N5S2.Rep.top | CAAACAT CATTACC CTCATTT CACCTAT AACTCT/3IAbRQSp/ |
| N5S2.Rep.bottom | /56-ROXN/TGAGTT ATAGGTG AAATGAG GGTAATG ATGTTTG AATGGTA |

#### Spurious displacement

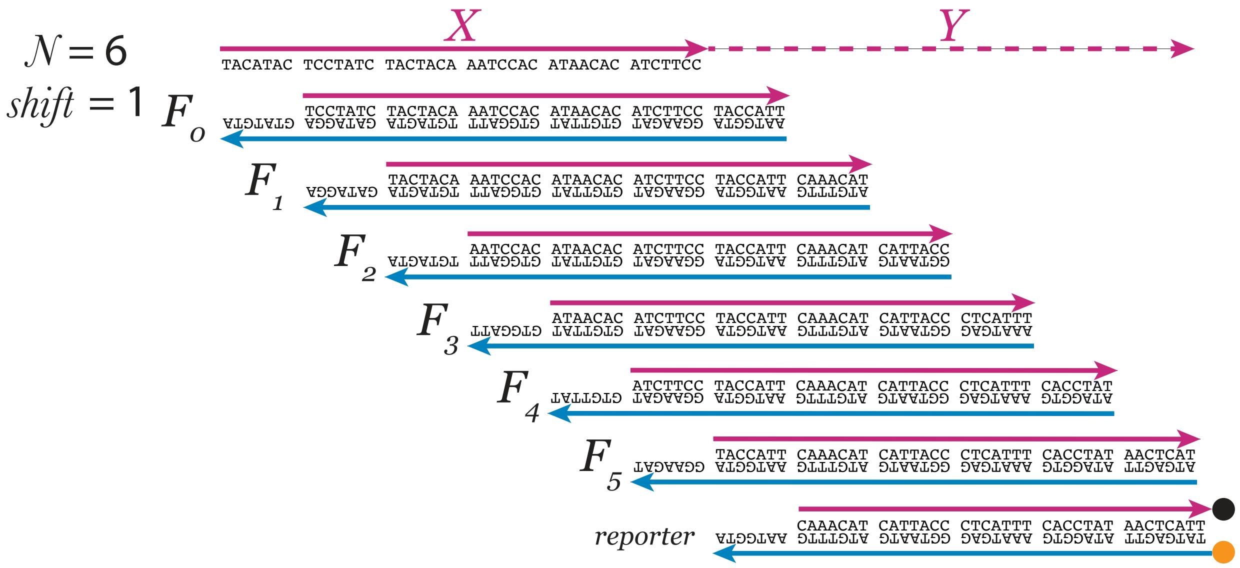

**Figure S7:** Sequences used for the experiment testing spurious displacement pathways in Figure 7.

| | |
|---|---|
| N6.X | TACATAC TCCTATC TACTACA AATCCAC ATAACAC ATCTTCC |
| N6S1.F0.top | TCCTATC TACTACA AATCCAC ATAACAC ATCTTCC TACCATT |
| N6S1.F0.bottom | AATGGTA GGAAGAT GTGTTAT GTGGATT TGTAGTA GATAGGA GTATGTA |
| N6S1.F1.top | TACTACA AATCCAC ATAACAC ATCTTCC TACCATT CAAACAT |
| N6S1.F1.bottom | ATGTTTG AATGGTA GGAAGAT GTGTTAT GTGGATT TGTAGTA GATAGGA |
| N6S1.F2.top | AATCCAC ATAACAC ATCTTCC TACCATT CAAACAT CATTACC |
| N6S1.F2.bottom | GGTAATG ATGTTTG AATGGTA GGAAGAT GTGTTAT GTGGATT TGTAGTA |
| N6S1.F3.top | ATAACAC ATCTTCC TACCATT CAAACAT CATTACC CTCATTT |
| N6S1.F3.bottom | AAATGAG GGTAATG ATGTTTG AATGGTA GGAAGAT GTGTTAT GTGGATT |
| N6S1.F4.top | ATCTTCC TACCATT CAAACAT CATTACC CTCATTT CACCTAT |
| N6S1.F4.bottom | ATAGGTG AAATGAG GGTAATG ATGTTTG AATGGTA GGAAGAT GTGTTAT |
| N6S1.F5.top | TACCATT CAAACAT CATTACC CTCATTT CACCTAT AACTCAT |
| N6S1.F5.bottom | ATGAGTT ATAGGTG AAATGAG GGTAATG ATGTTTG AATGGTA GGAAGAT |
| N6.rep.top.short | CAAACAT CATTACC CTCATTT CACCTAT AACTCATT/3IAbRQSp/ |
| N6.rep.bottom.short | /56-ROXN/TATGAGTT ATAGGTG AAATGAG GGTAATG ATGTTTG AATGGTA |

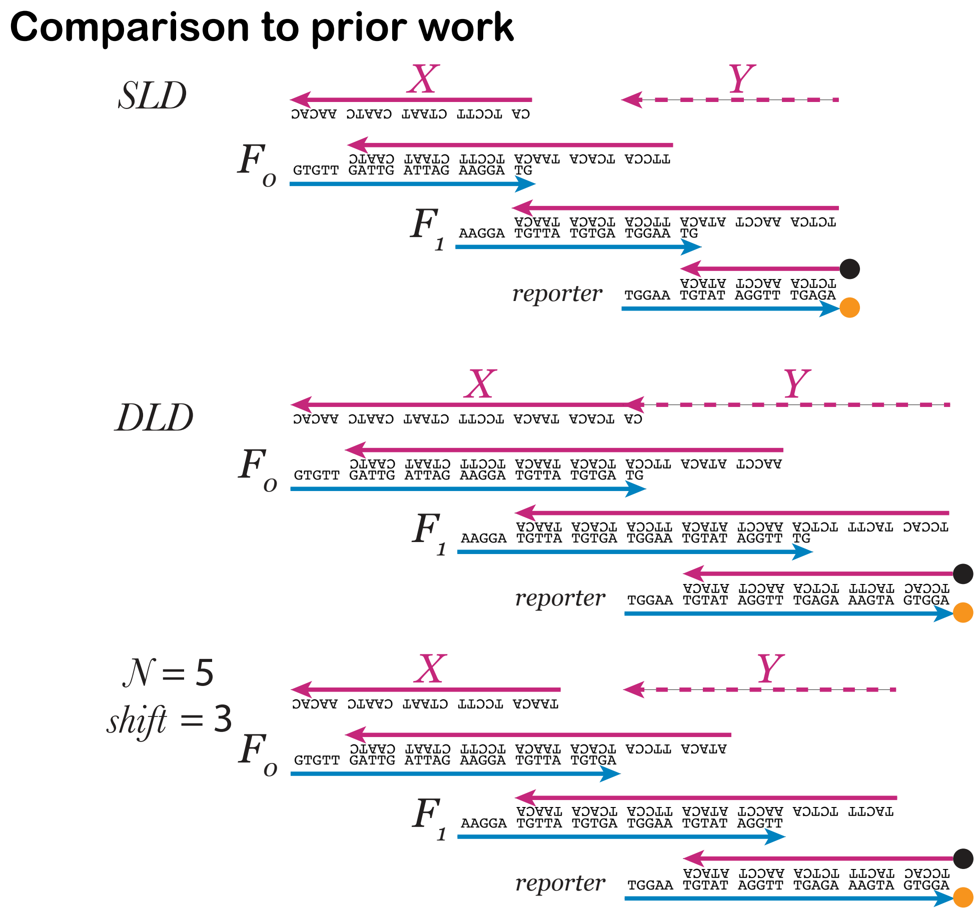

**Figure S8:** Sequences used for the experiment comparing the leak between the *SLD*, *DLD* in the prior work [3] and enthalpy-neutral DNA strand displacement design in Figure 8.

| | |
|---|---|
| SLD.F0.top | TTCCA TCACA TAACA TCCTT CTAAT CAATC |
| SLD.F0.bottom | GTGTT GATTG ATTAG AAGGA TG |
| SLD.F1.top | TCTCA AACCT ATACA TTCCA TCACA TAACA |
| SLD.F1.bottom | AAGGA TGTTA TGTGA TGGAA TG |
| SLD.Rep.top | /5IAbRQ/TCTCA AACCT ATACA |
| SLD.Rep.bottom | TGGAA TGTAT AGGTT TGAGA/3Rox_N/ |
| DLD.F0.top | AACCT ATACA TTCCA TCACA TAACA TCCTT CTAAT CAATC |
| DLD.F0.bottom | GTGTT GATTG ATTAG AAGGA TGTTA TGTGA TG |
| DLD.F1.top | TCCAC TACTT TCTCA AACCT ATACA TTCCA TCACA TAACA |
| DLD.F1.bottom | AAGGA TGTTA TGTGA TGGAA TGTAT AGGTT TG |
| DLD.Rep.top | /5IAbRQ/TCCAC TACTT TCTCA AACCT ATACA |
| DLD.Rep.bottom | TGGAA TGTAT AGGTT TGAGA AAGTA GTGGA/3Rox_N/ |
| N5.X | TAACA TCCTT CTAAT CAATC AACAC |
| N5.F0.top | ATACA TTCCA TCACA TAACA TCCTT CTAAT CAATC |
| N5.F0.bottom | GTGTT GATTG ATTAG AAGGA TGTTA TGTGA |
| N5.F1.top | TACTT TCTCA AACCT ATACA TTCCA TCACA TAACA |
| N5.F1.bottom | AAGGA TGTTA TGTGA TGGAA TGTAT AGGTT |
| N5.Reporter | share the same sequence as the DLD reporter |

#### Long cascade

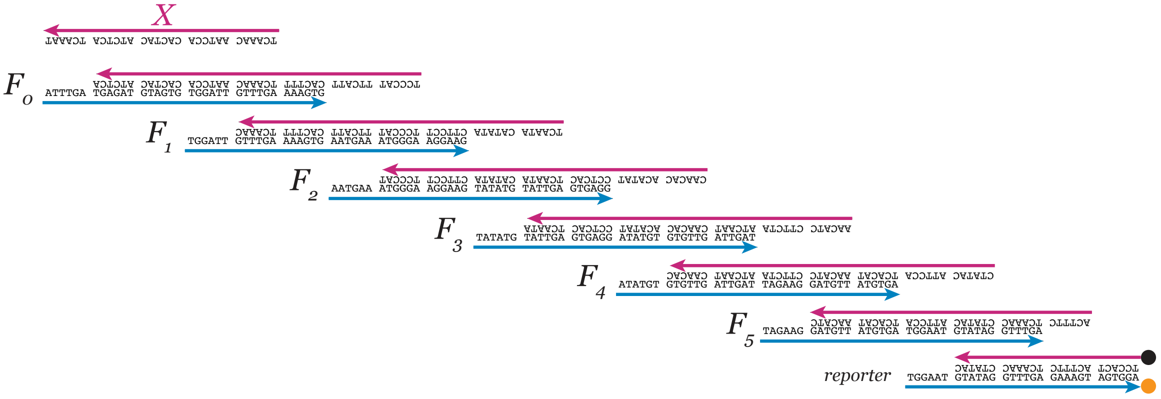

**Figure S9:** Sequences used for the long cascade experiment in Figure 9.

| | |
|---|---|
| Cascade.X0 | TCAAAC AATCCA CACTAC ATCTCA TCAAAT |
| Cascade.F0.top | TCCCAT TTCATT CACTTT TCAAAC AATCCA CACTAC ATCTCA |
| Cascade.F0.bottom | ATTTGA TGAGAT GTAGTG TGGATT GTTTGA AAAGTG |
| Cascade.F1.top | TCAATA CATATA CTTCCT TCCCAT TTCATT CACTTT TCAAAC |
| Cascade.F1.bottom | TGGATT GTTTGA AAAGTG AATGAA ATGGGA AGGAAG |
| Cascade.F2.top | CAACAC ACATAT CCTCAC TCAATA CATATA CTTCCT TCCCAT |
| Cascade.F2.bottom | AATGAA ATGGGA AGGAAG TATATG TATTGA GTGAGG |
| Cascade.F3.top | AACATC CTTCTA ATCAAT CAACAC ACATAT CCTCAC TCAATA |
| Cascade.F3.bottom | TATATG TATTGA GTGAGG ATATGT GTGTTG ATTGAT |
| Cascade.F4.top | CTATAC ATTCCA TCACAT AACATC CTTCTA ATCAAT CAACAC |
| Cascade.F4.bottom | ATATGT GTGTTG ATTGAT TAGAAG GATGTT ATGTGA |
| Cascade.F5.top | ACTTTC TCAAAC CTATAC ATTCCA TCACAT AACATC |
| Cascade.F5.bottom | TAGAAG GATGTT ATGTGA TGGAAT GTATAG GTTTGA |
| Cascade.rep.top | /5IAbRQ/TCCACT ACTTTC TCAAAC CTATAC |
| Cascade.rep.bottom | TGGAAT GTATAG GTTTGA GAAAGT AGTGGA/3Rox_N/ |
